# IL1 blockade attenuates *E. coli* induced intrauterine inflammation and preterm labor in Rhesus macaques

**DOI:** 10.64898/2026.07.30.741586

**Authors:** Pietro Presicce, Daniel Short, Monica Cappelletti, Natalia Perecki, Arne Gehlhaar, Alzbeta Godarova, Angela DeTomaso, Jacqueline Shauh, Liza Konnikova, Sivan Bercovici, Shiv Kale, Paul Babb, Lisa A. Miller, Sing Sing Way, William J. Zacharias, Hitesh Deshmukh, Senad Divanovic, Myung S. Sim, Alan H. Jobe, Claire A. Chougnet, Mark R. Johnson, Sam Mesiano, Suhas G. Kallapur

**Author notes:** **Correspondence:** Suhas G. Kallapur MD, Chief, Divisions of Neonatology and Developmental Biology, Professor of Pediatrics, David Geffen School of Medicine at UCLA, Mattel Children’s Hospital UCLA, 10833 Le Conte Avenue, Room B2-375 MDCC, Los Angeles, CA 90095, Email id. Equal contribution (Pietro Presicce, Daniel Short).

## Abstract

Intrauterine infection and inflammation are major causes of preterm labor, yet antibiotics alone often fail to prevent preterm birth despite achieving microbial clearance. Interleukin1 (IL1) is a key mediator of intra-amniotic inflammation and functional progesterone withdrawal, processes linked to labor initiation. We tested whether IL1 blockade with the clinically approved IL1 receptor antagonist, anakinra, could reduce inflammation and preterm labor in a Rhesus macaque model of intra-amniotic *Escherichia coli* infection followed by delayed antibiotic treatment. Here we show that anakinra reduced the incidence of preterm labor by 40% (p=0.07), attenuated inflammation in the fetal membranes and cervix, and preserved progesterone receptor-B abundance in the fetal membranes. These effects were most pronounced in animals protected from preterm labor. Our findings in a relevant model support the translational potential of combining IL1 blockade with antibiotics for infection-associated preterm labor.

**One sentence summary:** Recombinant human IL1 receptor antagonist (anakinra) decreased *E. coli* infection induced preterm labor and partially decreased intrauterine inflammation in a non-human primate model of inflammation mediated preterm labor.

## INTRODUCTION

Intrauterine infection (IUI) is a leading cause of preterm labor, a major public health concern, accounting for ∼40% of preterm births (PTB) [1, 2]. Additionally, IUI can also cause fetal inflammation or fetal inflammatory response syndrome (FIRS), a well-documented risk factor for adverse neonatal outcomes [3]. The source of microorganisms causing IUI is usually the lower genitourinary tract [4]. Although microbial invasion of the amniotic fluid is a known risk factor for PTB [5, 6], amniotic fluid cultures are rarely positive despite substantial intra-amniotic inflammation [5, 7]. Thus, the pathogenesis of microbe induced inflammation resulting in PTB remains a conundrum in the field.

Effective therapies for infection-associated preterm birth remain limited, as antibiotic treatment has generally failed to reduce the incidence of preterm birth in women with IUI [8], and in Rhesus macaque model of intrauterine infection [9]. These findings suggest that downstream inflammatory pathways may be important mediators. This concept shifts the exclusive therapeutic focus from pathogen eradication alone toward adjunctive targeted modulation of the inflammatory cascade.

IL1 was selected as a therapeutic target for this study based on prior studies implicating IL1 as a mediator of inflammation-induced PTBs [10–13] and IL1 is upstream of IL6 in the intrauterine inflammatory cascade [14]. This is particularly relevant because elevated IL6 is a robust biomarker of inflammation-associated PTB in both experimental models and human pregnancy [5, 7]. In addition, IL1 promotes functional progesterone withdrawal at the maternal-fetal interface by disrupting progesterone signaling, a pathway essential for the maintenance of pregnancy [15, 16].

To evaluate whether targeting the IL1 pathway can interrupt the inflammatory cascade leading to preterm labor, we used a Rhesus macaque model of intra-amniotic infection. To enhance clinical relevance, a uropathogenic strain of *E. coli* (UTI89) was used followed by delayed antibiotic treatment resulting in persistant inflammation despite antimicrobial therapy [9]. Because conventional microbial cultures frequently yield negative results despite clinical evidence of intra-amniotic infection, culture-based testing was supplemented with a highly sensitive, clinically validated assay for microbial cell-free DNA (cfDNA) detection [17]. To define the effects of IL1 blockade on the temporal dynamics of inflammation during infection-induced preterm labor, we longitudinally quantified key cytokines and neutrophil influx in the amniotic fluid and cervico-vaginal lavage, compartments previously implicated in the pathogenesis of preterm birth [18]. To assess tissue-level inflammatory responses, cytokine mRNA expression and neutrophil infiltration were evaluated in the fetal membranes and cervix at necropsy, as these gestational tissues are central mediators of labor-associated inflammation [19]. Given the essential role of progesterone signaling in maintaining pregnancy, we determined whether IL1 blockade preserves progesterone signaling in the setting of intrauterine inflammation [15, 16, 20].

## RESULTS

### Efficacy of anakinra for the prevention of preterm labor

Pregnant Rhesus macaques (∼85% gestation) were assigned to one of three groups (Fig. 1A): (i) IA saline/LB broth (control); (ii) IA inoculation with 10^6 colony-forming units (CFU) of live *Escherichia coli* followed by delayed antibiotic treatment initiated 24 hours later; or (iii) IA *E. coli* followed 24 hours later by antibiotics plus the IL-1 receptor antagonist anakinra, administered intra-amniotically and by maternal subcutaneous injection.The microbicidal antibiotic combination of cefazolin and enrofloxacin previously shown to have efficacy in this model were used [9]. Relatively large sample size for controls was due to adding tissues collected for previous studies [9]. There were no maternal/fetal deaths. Preterm labor (PTL) was defined by daily ultrasound and manual speculum examination of progressive cervical dilatation, shortening, and softening similar to clinical evaluation [21] (Fig. S1). Control animals did not experience PTL while 9/10 animals infected with *E. coli* showed definitive signs of PTL within 4 days (Fig. 1B). Anakinra-treated animals had a lower incidence of PTL compared with *E. coli*-exposed animals receiving antibiotics alone (55% [6/11] vs. 90% [9/11]), corresponding to a relative risk of 0.61 (relative risk reduction ∼40%); however, this difference did not achieve conventional statistical significance (p=0.07 Chi-square) (Fig. 1B). None of the animals had rupture of membranes prior to onset of labor. Of note, several PTL cases had vaginal births confirming bonafide preterm parturition (Table S1), while others had surgical delivery by C-section after diagnosis of progressive cervical changes to enable tissue harvest for assessments. To assess the probability of benefit within the constraints of limited non-human primate (NHP) sample size, we performed Bayesian Monte Carlo simulations (Fig. 1C). Using uninformative priors, the posterior difference in preterm labor rates between anakinra-treated (Group 3) and untreated animals (Group 2) had a 95% credible interval of −0.048 to 0.611, corresponding to a 96% posterior probability that anakinra reduced PTL incidence relative to antibiotics alone. There was some uncertainty of benefit as the credible interval crossed “0”.

**Fig. 1.**
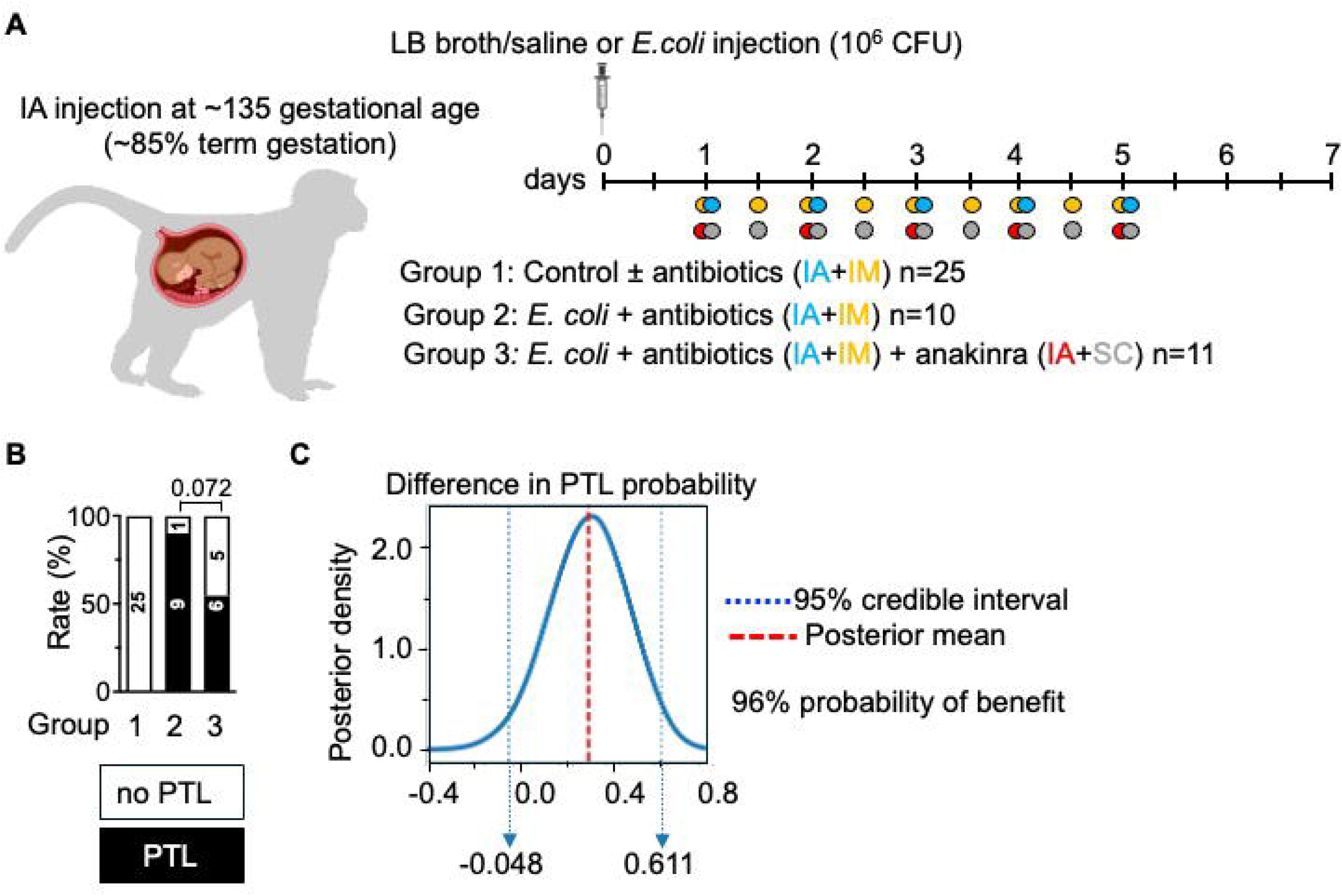
Effects of anakinra on preventing preterm labor. (**A**) Experimental model. Three groups of pregnant Rhesus macaque (*Macaca mulatta*) at ∼135d gestational age (∼85% term gestation) were given: 1) Intraamniotic (IA) saline/LB broth (Control n=25, with or without antibiotics (Abx)); 2) IA live *E. coli* (10^6^ CFU) followed one day later by Abx (n=10); 3) IA live *E. coli* (10^6^ CFU) followed one day later by Abx and anakinra (n=11). Abx regimen was as follows: IA cefazolin (10mg once daily) + IA enrofloaxicin (1mg once daily) + intramuscular (IM) cefazolin (25 mg/k twice daily) + IM enrofloaxicin (5mg/kg twice daily). Anakinra was administered IA (50mg/daily) + sub-cutaneous (SC) (100mg/2Xdaily). (**B**) Preterm labor (PTL) induced by *E. coli* was reduced from 9/10 to 6/11 after anakinra (p=0.072, chi-square t test), with no PTL in controls. (**C**) Bayesian Monte Carlo simulation with uninformative prior assumptions shows a 96% posterior probability of benefit with some uncertainty (95% credible interval crosses 0).

To understand if the difference in PTL outcome depended on drug pharmacokinetic variables, anakinra levels were measured longitudinally in the amniotic fluid (Fig. S2). Peak anakinra levels were reached 24h after administration in the amniotic fluid with values 7.2 ± 8.2 µg/mL (mean ± SD). Importantly, no differences in levels were noted between groups experiencing preterm labor vs. those protected, suggesting that variations in anakinra drug concentrations do not explain differences in PTL outcome.

### Efficacy of antibiotics and measurement of microbial products

To understand the relationship between preterm labor and the burden of infection, amniotic fluid (AF) and maternal blood were sampled daily for bacterial cultures. IA injection of live *E. coli* led to a robust growth with positive AF culture 24h after *E. coli* injection and prior to starting antibiotics (Fig. 2A). The antibiotic regimen was highly effective, resulting in elimination of *E. coli* in all injected animals starting 24h after antibiotics with durable effects persisting until delivery (Fig. 2A). Importantly, inhibition of IL1 signaling by anakinra did not alter bactericidal efficacy of antibiotics. Notably, even when AF cultures showed robust *E. coli* growth 24h after injection, the matched maternal blood cultures demonstrated no growth. Similarly, none of the cord blood samples or fetal liver cultures grew *E. coli* (Fig. 2B). As expected, none of the control animals had *E. coli* growth in AF or other body fluids at any time point (Fig. 2A-B). Despite the absence of cultivatable organisms at delivery in the *E. coli*–exposed group, *E. coli* cell-free DNA (cfDNA) levels in amniotic fluid increased by approximately five orders of magnitude from pre-injection baseline levels to delivery (Fig. 2C). Anakinra did not significantly change *E. coli* cfDNA titers (Fig. 2C). Interestingly, *E. coli* cfDNA titers in the AF at delivery were generally higher in animals that developed PTL compared to those with no PTL (Fig. 2D). In contrast to the AF, maternal plasma *E. coli* cfDNA increased only modestly between pre-exposure to delivery (Fig. 2E). Although cord blood cultures showed no *E. coli* growth, *E. coli* cfDNA was readily detectable (Fig. 2F). Comparison of different compartments within the same subject revealed that *E. coli* cfDNA titers at delivery were AF>>cord blood>maternal blood (Fig. 2F), indicating distinct compartmentalization and selective trafficking of microbial cfDNA. Similar to the AF, the fetus with lowest *E. coli* cfDNA was not born after PTL (Fig. 2F). The generally lower AF and cord blood *E. coli* cfDNA in animals with no PTL regardless of anakinra treatment suggest a potential association between microbial cfDNA burden and PTL.

**Fig. 2.**
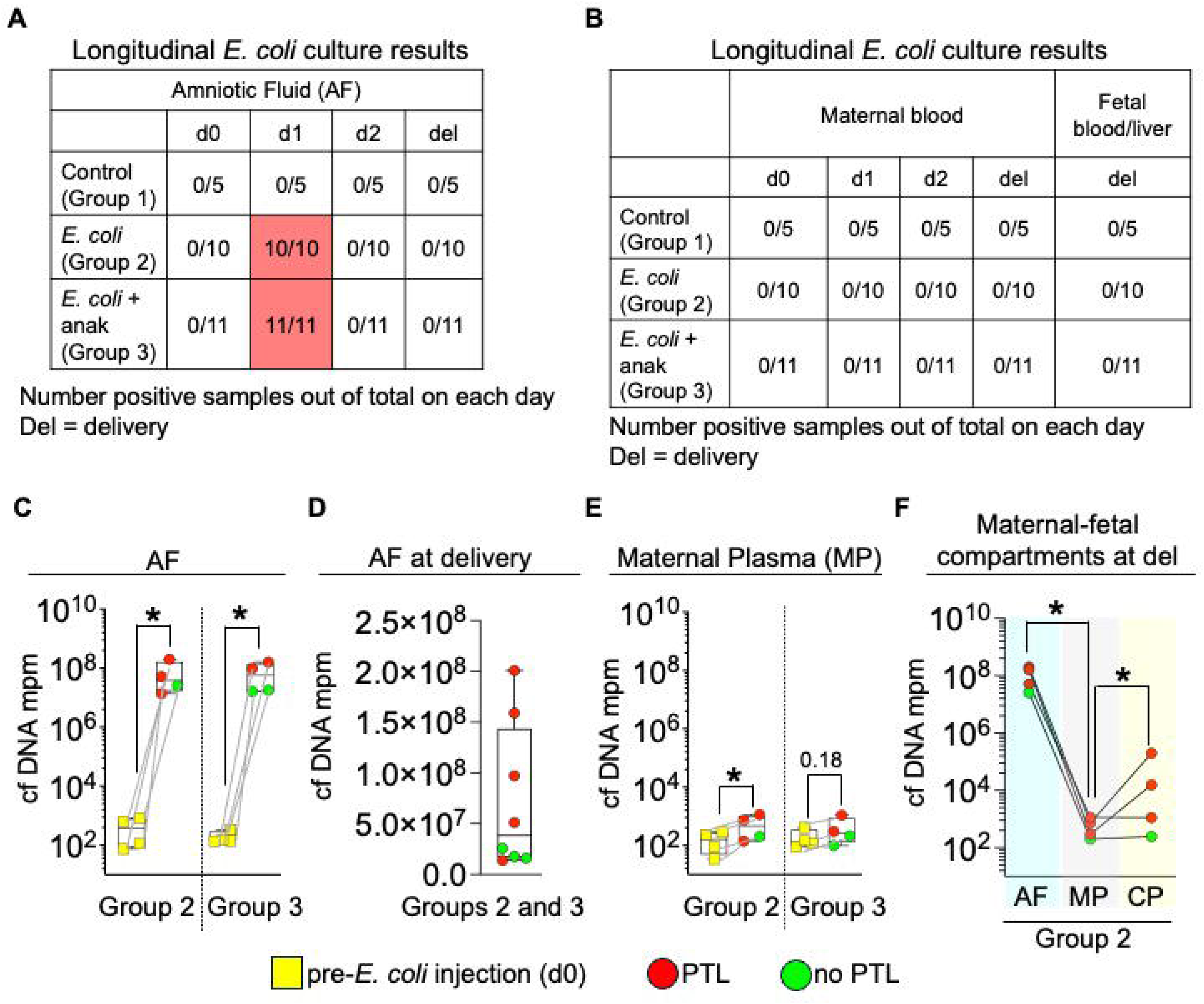
Antibiotics efficiently clear bacteremia, but *E. coli* cell free DNA persists. (**A-B**) Amniotic fluid (AF) and maternal blood were collected daily and cord blood/fetal liver were collected at necropsy. *E. coli* growth at different time points (day 0=pre-exposure; del=delivery) is shown. Note that all *E. coli* injected animals were culture positive in the AF at 24h but negative at delivery. Maternal or fetal blood cultures were negative at all times tested (**C**) *E. coli* cell free (cf) DNA titers depicted as molecules per microliter (MPM) significantly increased in the AF from pre-exposure to delivery in *E. coli*-exposed animals with no effect of anakinra. Comparison is in the same animal from pre-*E. coli* injection and delivery (green dot =no PTL, red dots =PTL). (**D**) Animals with PTL showed generally higher *E. coli* cfDNA titers in the AF compared to those with no PTL. (**E**) Changes from pre-exposure to delivery were modest in maternal plasma. (**F**) *E. coli* cfDNA in the different maternal-fetal compartments within the same pregnancy suggesting more selective trafficking of AF cfDNA to fetal compared to maternal circulation (AF=amniotic fluid; MP=maternal plasma; CP=cord plasma). Mann–Whitney U-test was used for two group comparison after stratification of the anakinra group by PTL status. *p < 0.05 between comparators is shown.

### Inflammation in reproductive tissues at delivery

To examine functional outcomes after *E. coli*, inflammatory markers in the fetal membranes (comprising the chorion, amnion, and decidua parietalis), the cervix, and myometrium were evaluated as these tissues are most affected by intrauterine infection [4, 19].

#### Fetal membranes

Compared to low levels in controls, *E. coli* caused large increases (several hundred-fold) in mRNAs for pro-inflammatory cytokines/chemokines *IL1B, IL6, CXCL8/IL8* and neutrophil frequency in the fetal membranes (Fig. 3A). Similar increases were also observed in mRNAs for *MCP1/CCL2, TNF,* and the gene for a key enzyme in prostaglandin synthetic pathway *PTGS2* (Fig. S3A). The entire anakinra group regardless of preterm labor outcome had significantly or nearly significantly decreased levels of *IL1B, IL6, CXCL8/IL8* and neutrophil frequency induced by *E. coli* (Fig. 3A), but anti-inflammatory effects of anakinra were less evident for *MCP1/CCL2*, *TNF* and *PTGS2* (Fig. S3A). Of note, the anakinra subgroup with no preterm labor had the lowest levels of inflammatory markers approaching control levels (Fig. 3A and S3A).

**Fig. 3.**
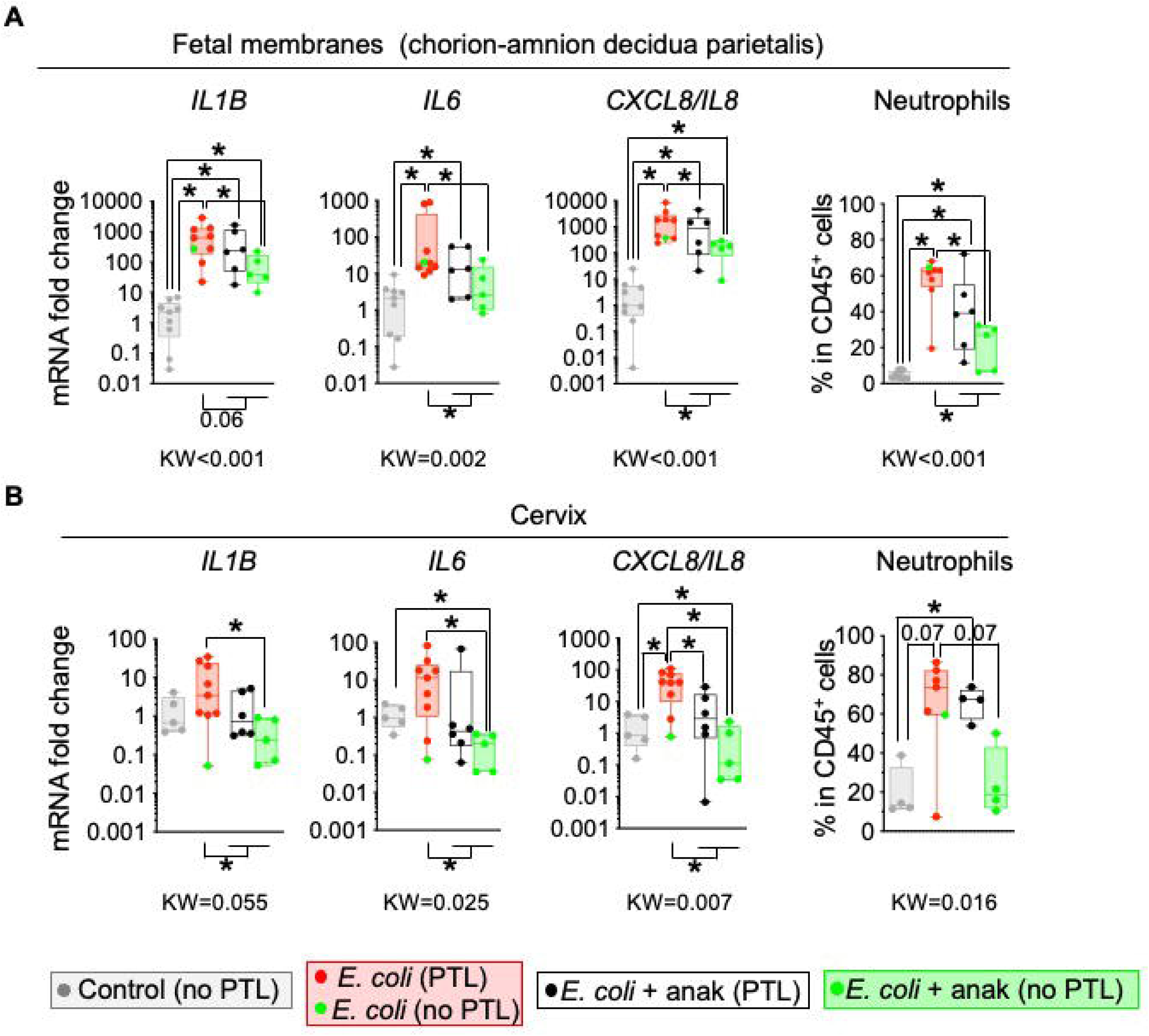
Anakinra reduced *E. coli* induced inflammatory markers in reproductive tissues at delivery. Extraplacental chorio-amnion decidua parietalis tissue (fetal membranes) and cervix were analyzed for cytokine mRNA expression by quantitative PCR using Taqman probes. Transcript abundance was normalized to 18S RNA and expressed as fold change relative to controls. Neutrophils were quantified by flow cytometry of cell suspensions. (**A**) In the fetal membranes, *E. coli* exposure markedly increased *IL1B*, *IL6*, *CXCL8/IL8 mRNAs* compared to controls. Anakinra decreased *E. coli* induced inflammatory markers with highest decreases in animals without preterm labor (PTL). (**B**) In the cervix, *E. coli* exposure increased *CXCL8/IL8* mRNA and neutrophil frequency had trends towards increase (p=0.07). Anakinra decreased *E. coli* induced *IL1B*, *IL6*, *CXCL8/IL8* mRNAs with highest decreases in animals without preterm labor. Data are mean ± SEM. Initial all group comparisons were performed using the Kruskal-Wallis (KW) test, followed by Mann–Whitney U-test for two group comparison after stratification of the anakinra group by PTL status. *p < 0.05 between comparators is shown.

#### Cervix

Since cervical ripening is an integral component of labor, inflammation was assessed in this tissue. Compared to controls, *E. coli* increased *CXCL8/IL8* mRNA with a trend towards increase for neutrophil proportions but mRNAs for *IL1B* and *IL6* did not increase significantly (Fig. 3B). Similarly, *E. coli* increased mRNAs for *PTGS2,* but not *MCP1/CCL2* or *TNF* (Fig. S3B). Similar to the fetal membranes, anakinra treatment regardless of PTL outcome significantly or nearly significantly decreased *E. coli* induced mRNAs for *IL1B, IL6, CXCL8/IL8* (Fig. 3B*), and MCP1/CCL2, TNF, PTGS2* (Fig. S3B). Notably, the subgroup without preterm labor had the lowest mRNA levels. Compared to controls, *E. coli* cervix neutrophil proportions were borderline significantly increased. Anakinra subgroup with no PTL had a trend towards decreased *E.coli* induced neutrophil proportions (p=0.07) (Fig. 3B).

#### Myometrium

We previously reported that intra-amniotic injection of LPS induces minimal inflammation in the myometrium [22]. Here we evaluated effects of exposure to intra-amniotic live organism. Compared to controls, *E. coli* increased mRNAs for pro-inflammatory cytokines/chemokines *IL1B, IL6, CXCL8/IL8, MCP1/CCL2*, *TNF,* and *PTGS2* (Fig. S3C). Anakinra did not significantly reduce *E. coli* induced pro-inflammatory gene expression regardless of the preterm labor outcome (Fig. S3C). In contrast to a massive neutrophil influx in the fetal membranes, *E. coli* with or without anakinra did not significantly change neutrophil proportions within the immune cells in the uterus Fig. S3C. These findings suggest that fetal membranes and the cervix more than the myometrium are targets of tissue level inflammation during intrauterine infection. Among the contraction associated-genes, *E. coli* increased mRNAs for *GJA1* (connexin 43) only, but not for *OXTR, PTGFR,* and *PTGS1* (Fig. S3D). Anakinra group had similar contraction associated-gene expression compared to the *E. coli* group (Fig. S3D).

### Temporal trajectories of inflammatory products in maternal compartments

Since tissue level inflammation was evaluated at delivery after the diagnosis of preterm labor, we asked if anakinra influenced inflammatory signals prior to the onset of labor. This was accomplished using longitudinally collected daily sampling of the amniotic fluid, cervico-vaginal lavage, and maternal blood.

#### Amniotic fluid (AF)

*E. coli* robustly increased AF levels of pro-inflammatory cytokines IL1*β*, IL6, CXCL8/IL8, and neutrophil counts with peak levels 24h after injection (Fig. 4A). Similar AF increases were also seen with TNFα, MCP1/CCL2, and GM-CSF (Fig. S4A). Compared to pro-inflammatory cytokines, the anti-inflammatory cytokine IL10 increased more modestly (Fig. S4A). Anakinra treated group had similar trajectories of changes in cytokines as the *E. coli* group regardless of preterm labor outcome (Fig. 4A and Fig. S4A). As expected, control animals without *E. coli* exposure had no significant change in AF cytokines and neutrophil counts from baseline (Fig. 4A and Fig. S4A).

**Fig. 4.**
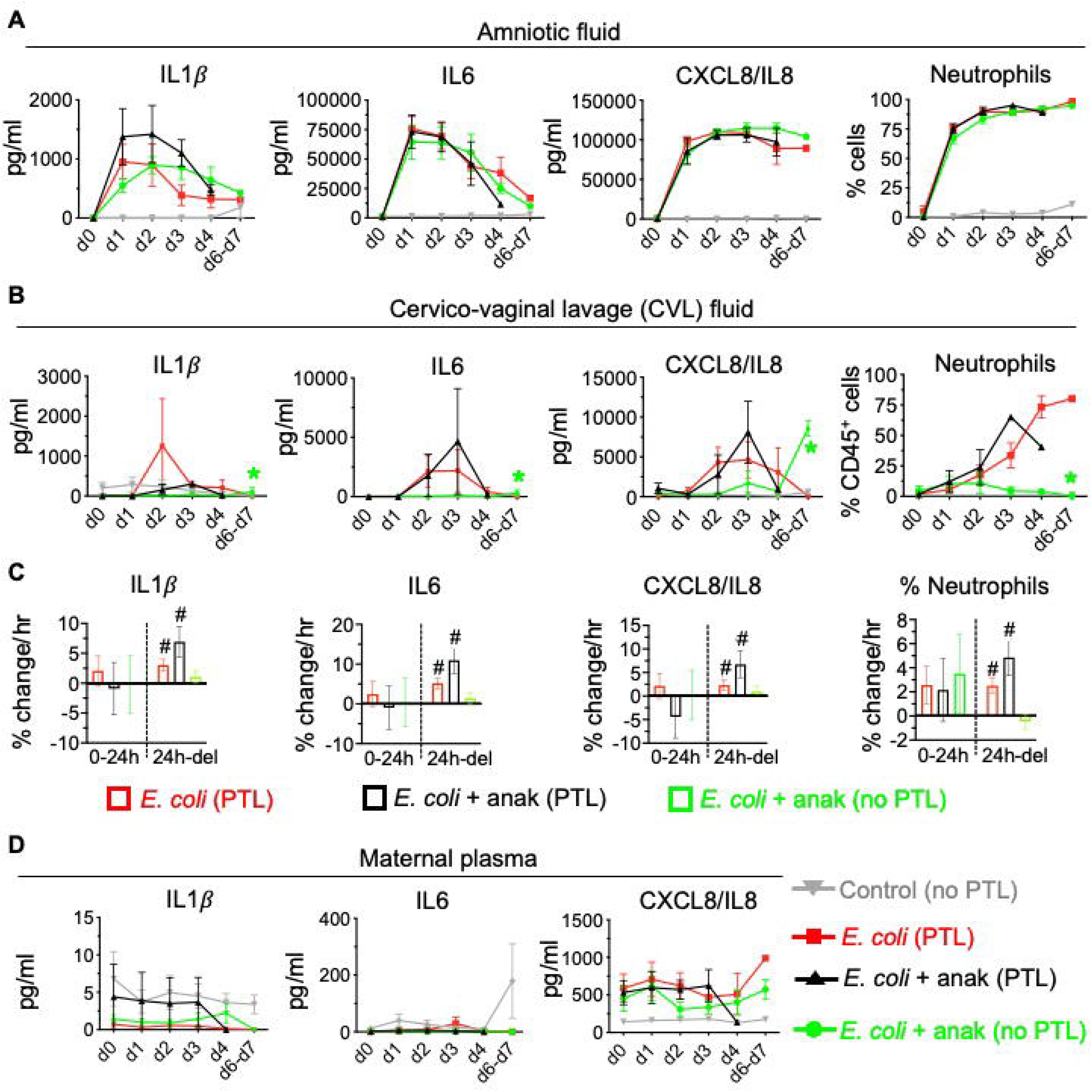
Inflammatory response trajectories to *E.coli* infection differed in different maternal compartments. Cytokine concentrations were measured longitudinally by multiplex immunoassay. Neutrophil frequencies were quantified by Diff-Quik staining of cytospin preparations in amniotic fluid and by flow cytometry in cervico-vaginal lavage (CVL) fluid. (**A**) Amniotic fluid (AF). Relative to controls, intra-amniotic *E. coli* exposure induced marked increases in cytokine concentrations and neutrophil frequencies. Temporal trajectories were similar among all *E. coli*–exposed groups and were not significantly affected by anakinra treatment or preterm labor outcome. (See Fig S4 for AF mixed-effect modeling data for inflammatory trajectories). (**B**) Cervico-vaginal lavage fluid (CVL). Compared with controls, *E. coli* exposure increased cytokine concentrations and neutrophil frequencies, with onset delayed relative to the AF compartment. Animals treated with anakinra that subsequently developed preterm labor exhibited inflammatory trajectories similar to those of the *E. coli* group. In contrast, animals treated with anakinra that did not develop preterm labor displayed attenuated inflammatory trajectories after day 1. Green asterisks indicate significant differences in trajectory from d1 to delivery between (*E. coli* and anakinra preterm labor) groups and anakinra-treated animals without preterm labor as determined by mixed-effects modeling. (**C**) Mixed-effects analysis of CVL inflammatory trajectories. Longitudinal trajectories were modeled in two prespecified epochs separated by treatment initiation at 24 hours after *E. coli* inoculation. During the first 24 hours, regression slopes were not significantly different from zero in any group. From 24 hours to delivery, the *E. coli* group and the anakinra-treated subgroup that developed preterm labor exhibited significant positive slopes (#, p<0.05 compared to zero value), whereas slopes in the anakinra-treated subgroup without preterm labor were not significantly different from zero. (**D**) Maternal plasma. Cytokine concentrations remained largely unchanged over time in control and *E. coli*–exposed animals, irrespective of treatment group.

#### Cervico-vaginal lavage fluid (CVL)

In contrast to the AF, CVL peak levels were achieved at d2-3, which is later and of lower magnitude compared to the AF (Compare Figs. 4A and B). *E. coli* increased CVL fluid levels of IL1*β*, IL6, CXCL8/IL8, and neutrophil counts (Fig. 4B). As expected, control animals without *E. coli* exposure had no significant change in CVL cytokines from baseline (Fig. 4B). Similar CVL increases were also seen with TNFα, MCP1/CCL2, GM-CSF and the anti-inflammatory cytokine IL10 (Fig. S5A). Interestingly, in the anakinra subgroup with no PTL a flat trajectory of cytokines was noted, similar to controls. This flat trajectory was in contrast with increasing inflammatory marker levels for those experiencing PTL (Fig. 4B and Fig. S5A).

#### Trajectory analyses

To determine whether inflammatory trajectories differed among treatment groups, we applied linear mixed-effects models to longitudinal cytokine and cellular measurements in AF and CVL. Fixed effects included treatment group, time, and treatment-by-time interaction terms and inter-animal biological variability were incorporated as random effects. To understand the effects of antibiotic ± anakinra treatment intervention starting at 24h, time variables were segmented separately for the periods 0-24h vs. 24h to delivery. Regression slopes (% change/hr) and treatment-by-time interactions were compared across groups.

The CVL cytokine concentrations remained largely unchanged during the first 24 hours after intra-amniotic *E. coli* exposure as noted by regression slope values near “0” for all groups (Fig. 4C and Fig. S5B). In contrast, for the period 24h onward, there was a divergent trajectory for those experiencing PTL vs. no PTL. In the *E. coli* group and the anakinra-treated subgroup that subsequently developed PTL, IL6, IL1β, CXCL8/IL8, MCP1/CCL2, GM-CSF, TNFα, and IL10 and neutrophil frequency continued to increase over time (positive regression slope values in Fig. 4C and Fig. S5B). In contrast, these inflammatory markers remained stable in anakinra-treated animals that did not develop PTL, with % change/hr slope values approaching zero. A notable exception to this trend was the anti-inflammatory cytokine IL10, whose levels continued to increase (regression slope was positive) after 24h in all groups (Fig. S5B). Notably, suppression of CVL inflammatory trajectories occurred before the onset of labor, indicating that successful IL1 blockade was associated with early arrest of the inflammatory cascade preceding preterm birth.

In the AF, IL1β, IL6, TNFα, CXCL8/IL8, MCP1/CCL2, GM-CSF, and IL10 regression slopes increased during the first 24 hours after intra-amniotic *E. coli* inoculation (positive values) and declined following antibiotic administration (negative values), with similar temporal patterns across treatment groups irrespective of anakinra exposure or preterm labor outcome (Fig. S4B). These results suggest that antibiotics were the primary driver of reductions in cytokine concentrations. In contrast, *E. coli*–induced elevations in AF CXCL8/IL8 and IL10 and neutrophil counts remained elevated until delivery despite antibiotic treatment (Fig. 4A and Fig. S4B).

#### Immune cell infiltration in the CVL

Because cervico-vaginal lavage (CVL) inflammation emerged as a key discriminator of preterm labor outcome, we next characterized the cellular composition of this compartment (Fig. S5C). Flow cytometry data visualized by t-distributed stochastic neighbor embedding (t-SNE) revealed a marked shift from a predominantly nonimmune cell population at baseline to a leukocyte-rich inflammatory infiltrate at delivery following intra-amniotic *E. coli* exposure. The infiltrate comprised multiple immune cell populations, including neutrophils, macrophages, T cells, B cells, NK cells, and NKT cells (Fig. S5C). Total CD45^+^ leukocyte counts increased by approximately 50-fold at delivery relative to pre-exposure levels, with neutrophils representing the dominant immune cell population across *E. coli*–exposed groups (Fig. S5D). Anakinra reduced overall CVL leukocyte accumulation regardless of preterm labor outcome (Fig. S5D). However, a reduction in neutrophil abundance was observed only in anakinra-treated animals that did not develop preterm labor (Fig. S5D), consistent with the attenuated cytokine trajectory observed in this subgroup. These findings identify suppression of neutrophil recruitment as a distinguishing feature of successful protection from inflammation-induced preterm labor.

#### Maternal blood

Maternal plasma concentrations of IL1β, IL6, CXCL8/IL8 (Fig. 4D), MCP1/CCL2, GM-CSF, TNFα, and IL10 (Fig. S6) did not increase after *E. coli* in sharp contrast with the trajectories in the AF and CVL. The absence of inflammatory mediator increases in maternal plasma is consistent with minimal increases in *E. coli* cfDNA in maternal blood (Fig. 2E). These data demonstrate that even after exposure to a virulent organism in the amniotic space, inflammation was contained in the intrauterine compartment without significant maternal systemic involvement, recapitulating clinical data demonstrating distinct compartmentalization and localized nature of intrauterine inflammation [23].

#### Anakinra pharmacokinetics

To understand if divergent anakinra anti-inflammatory responses were due to different local drug concentrations, anakinra levels were measured in the CVL. Anakinra levels were ∼3-logs lower compared to the amniotic fluid, with peak levels values0.006422 ± 0.00665 µg/ml (mean ± SD) (Fig. S2). Importantly and similar to the amniotic fluid, no differences in anakinra levels were noted between groups with and without preterm labor or with different inflammatory marker trajectories.

### Progesterone signaling in the extraplacental fetal membranes

Progesterone receptor (PR) signaling, particularly through the transcriptionally active PR-B isoform, is essential for the maintenance of pregnancy [24, 25]. PR-B differs from the inhibitory PR-A isoform by an additional 164 amino acids at its N-terminus, enabling selective detection with a PR-B–specific antibody. While prior studies quantified total PR expression using antibodies that recognize both PR-A and PR-B [26], we specifically assessed PR-B abundance in gestational tissues. In control animals, robust nuclear PR-B immunostaining was observed in cells morphologically consistent with decidual stromal cells at the maternal-fetal interface identifying the decidual stroma cells as key targets for progesterone activity (Fig. 5A). Intra-amniotic *E. coli* exposure significantly reduced nuclear PR-B immunostaining in decidual stromal cells (Fig. 5A, C). Although PR-B was not significantly different between the *E. coli* and the overall anakinra-treated groups, animals treated with anakinra that did not develop preterm labor exhibited significantly greater PR-B immunostaining in decidual stromal cells than animals exposed to *E. coli* alone (Fig. 5C).

**Fig. 5.**
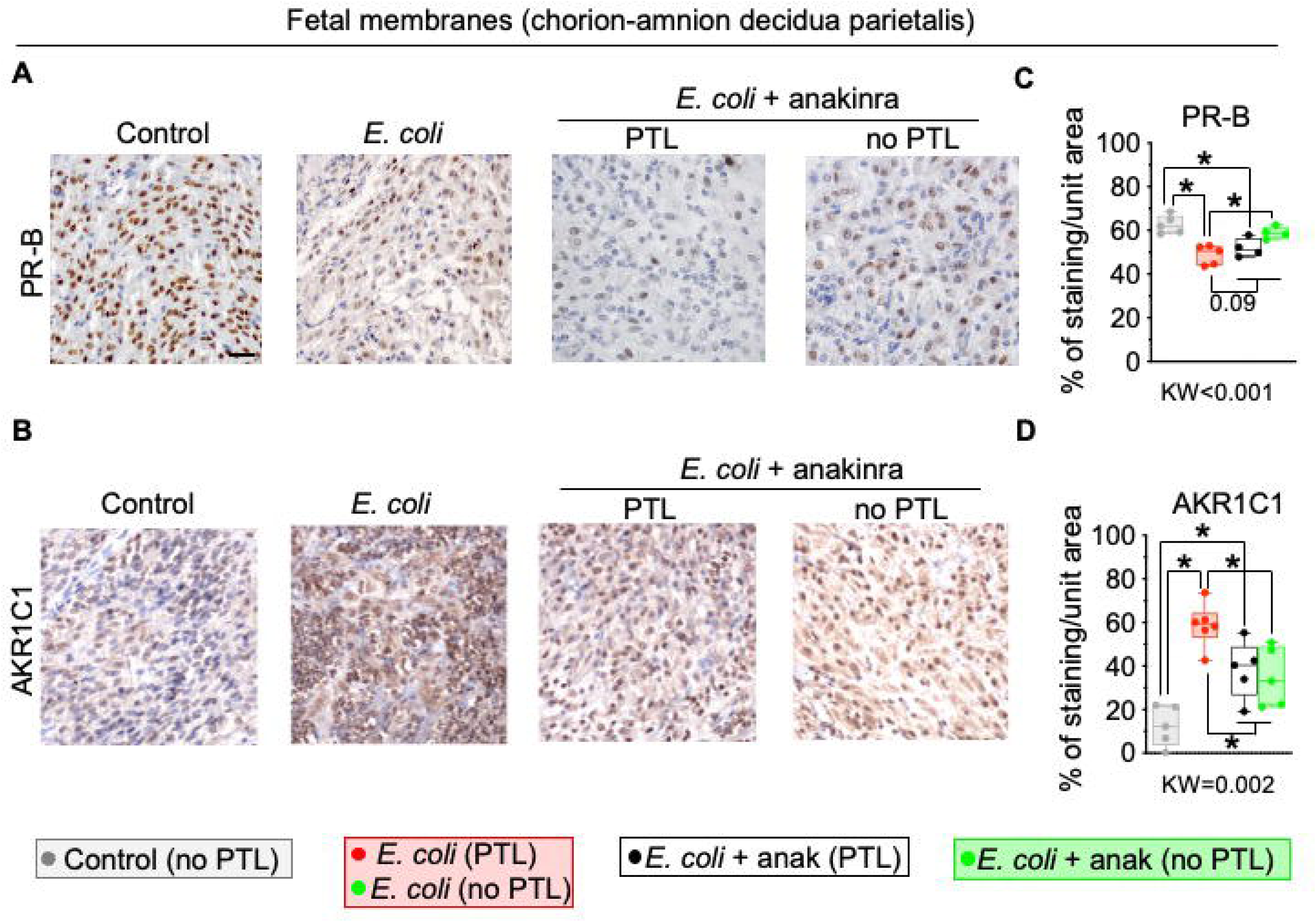
Anakinra partially restored *E. coli* induced disruptions in progesterone receptor signaling in the fetal membranes. Representative images showing immunohistological detection (brown staining) of (**A**) Progesterone receptor-B isoform (PR-B) (n=4-5/group) and (**B**) AKR1C1 (n=5-6/group) in cells with morphologic appearance of decidua stroma cells. Quantitative analysis showed that (**C**) *E. coli* exposure decreased PR-B^+^ cells. The anakinra treated group showed dichotomous outcomes: animals with preterm labor (PTL) had similar reduction in PR-B abundance, while the no PTL group had partial restoration of PR-B expression (**D**) *E. coli* exposure significantly increased AKR1C1^+^ cells while the anakinra treated animals with no PTL showed decreased AKR1C1+ cells compared to *E. coli* animals. Data are mean ± SEM. Initial all group comparisons were performed using the Kruskal-Wallis (KW) test, followed by Mann–Whitney U-test for two group comparison after stratification of the anakinra group by PTL status. *p < 0.05 between comparators is shown. (scale bar = 20 μm).

Inflammation-induced functional progesterone withdrawal is also mediated by increased expression of aldo-keto reductase family 1 member C1 (AKR1C1), a progesterone-inactivating enzyme expressed by decidual stromal cells [16]. *E. coli* exposure increased AKR1C1 immunostaining in the fetal membranes, whereas anakinra reduced AKR1C1 expression regardless of preterm labor outcome (Fig. 5 B, D). Together, these findings indicate that IL1 blockade preserves key components of progesterone signaling (i.e., PR-B and progesterone) in inflamed gestational tissues.

### Fetal inflammation

A major consequence of intrauterine infection is fetal inflammatory response syndrome (FIRS), which results from fetal exposure to inflammatory mediators within the amniotic cavity and is commonly defined by elevated cord blood IL6 concentrations [3]. Consistent with the development of FIRS, intra-amniotic *E. coli* exposure significantly increased cord blood IL6 and CXCL8/IL8 but not IL1β and CCL2/MCP1 concentrations compared with controls (Fig. 6A). Although cytokine concentrations in the overall anakinra-treated group were similar to those in the *E. coli* group, animals treated with anakinra that did not develop preterm labor exhibited significantly lower cord blood IL6 and CCL2/MCP1 concentrations than animals exposed to *E. coli* alone (Fig. 6A), indicating attenuation of the fetal systemic inflammatory response in this subgroup.

**Fig. 6.**
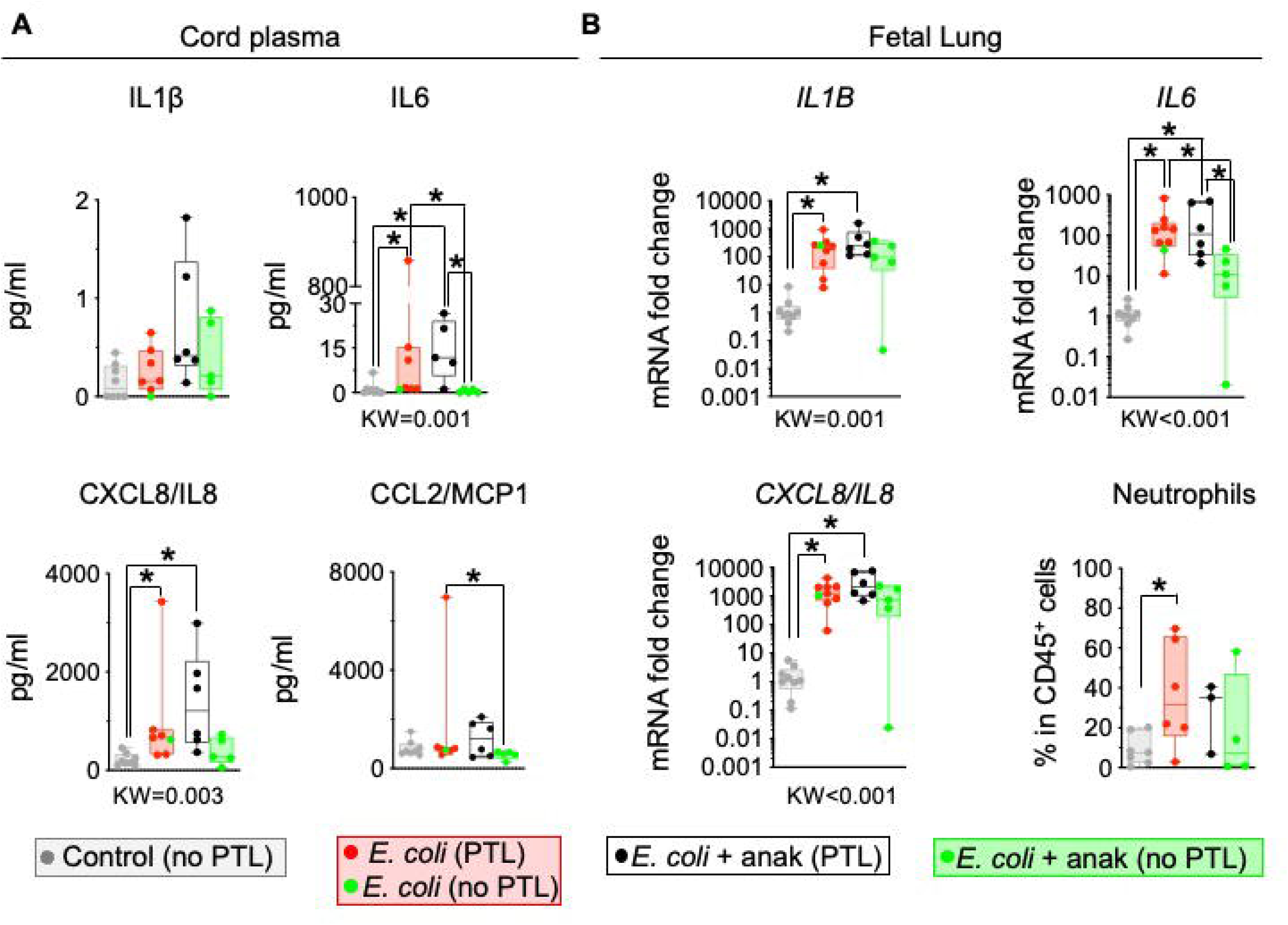
Anakinra partially decreased *E. coli* induced fetal inflammation. (**A**) Plasma from umbilical cord blood at delivery was used for measuring cytokine protein concentrations by multiplex assay. Cord blood IL6 and CXCL8/IL8 increased in the *E. coli* group compared to controls indicating fetal inflammatory response syndrome. In the anakinra subgroup with no PTL, cord blood IL6 and CCL2/MCP1 levels were lower compared to the *E. coli* group. (**B**) Fetal lung cytokine mRNA expression was evaluated by quantitative PCR using Taqman probes. Transcript abundance was normalized to 18S RNA and expressed as fold change relative to controls. Neutrophils were quantified by flow cytometry of dissociated fetal lung cell suspensions. *E. coli* and anakinra subgroup with PTL increased *IL1B, IL6, IL8/CXCL8* mRNAs, and neutrophil frequencies. Anakinra subgroup without PTL had lower *E. coli* induced *IL6* mRNA. Data are mean ± SEM. Initial all group comparisons were performed using the Kruskal-Wallis (KW) test, followed by Mann–Whitney U-test for two group comparison after stratification of the anakinra group by PTL status. *p < 0.05 between comparators is shown.

The fetal lung is particularly susceptible to intrauterine inflammation because fetal breathing movements continuously expose the developing lung to inflammatory mediators present in the amniotic fluid [4]. Accordingly, *E. coli* exposure markedly increased pulmonary expression of the pro-inflammatory cytokines and chemokines mRNAs for *IL1B, IL6, CXCL8/IL8* and promoted neutrophil infiltration into the fetal lung (Fig. 6B). Similar increases were also seen in mRNAs for *CCL2/MP1, TNF,* and *PTGS2* (Fig. S7). While the overall anakinra-treated group did not show significant reductions in pulmonary inflammatory gene expression compared with the *E. coli* group, animals treated with anakinra that remained protected from preterm labor exhibited significantly lower *IL6* mRNA expression (Fig. 6B and Fig. S7). In addition, neutrophil infiltration was more modest following anakinra treatment, since neutrophil frequencies did not increase significantly compared to control values (Fig. 6B). Together, these findings suggest that successful IL1 blockade mitigates some aspects of fetal systemic and fetal lung inflammation in the subgroup that also is protected from preterm labor.

## DISCUSSION

Preterm birth affecting ∼15 million births/yr globally remains a major unmet clinical challenge, with no approved therapies to prevent inflammation-driven labor [27]. A strength of this study is the use of a rigorously characterized non-human primate model in which delayed antibiotic treatment closely recapitulates the clinical scenario of intrauterine infection–associated preterm labor. Inhibition of IL1 signaling by anakinra, a clinically approved drug, partially reduced *E. coli* induced inflammation at the maternal-fetal interface with a ∼40% relative risk reduction in the rate of preterm labor. Anti-inflammatory effects of anakinra were most pronounced in the extraplacental fetal membranes and the cervix. The attenuation of inflammation was the greatest in the subgroup with no preterm labor. Indeed, in this benefited subgroup, post–*E. coli* induced cytokine increases in the cervico-vaginal lavage fluid were arrested in animals before the onset of preterm labor. The observed association between inflammatory burden and preterm labor is consistent with the concept of an “inflammatory threshold” for its induction. Another important finding was that IL1 blockade by anakinra partially reversed the decrease in pregnancy-sustaining PR-B abundance in decidual stromal cells and prevented the local inactivation of progesterone by AKR1C1. Together, these changes would boost the progesterone/PR-B signal maintaining the protective effect of progesterone to block parturition. This pre-clinical study provides mechanistic clues of how IL1 receptor antagonist therapy may be beneficial in treating inflammation mediated preterm labor.

Given that exposure to live *E. coli* was transient due to rapid antibiotic efficacy, what mechanisms account for the persistence of inflammatory responses? One possible explanation is the *E. coli* cell-free DNA (cf DNA) concentrations which increased by ∼5-log orders in the amniotic fluid. The concordance between microbial cf DNA titers and the magnitude of inflammation in the compartments AF>>cord blood>maternal blood suggests that microbial cfDNA is a candidate pathogen associated molecular pattern (PAMP) triggering intrauterine inflammation. Our experiments and previous studies suggest that amniotic microbial cfDNA is more readily trafficked to fetal blood compared to maternal blood (Fig. 2F) [28]. A central paradox in the field is the high prevalence of cases with negative amniotic fluid cultures despite substantial intra-amniotic inflammation [5, 7]. Together with prior studies [28], our findings suggest a potential explanation for this discrepancy. We propose that ascending microorganisms may localize within the chorio-decidua while releasing microbial products into the amniotic cavity. These products, including microbial cf DNA acting as pathogen associated molecular patterns (PAMPs) [29, 30], can be detected by the fetal and maternal innate immune system through multiple nucleic acid–sensing pathways, including TLR-mediated recognition of hypomethylated CpG motifs, cGAS– STING signaling, and AIM2 inflammasome activation [31]. Activation of these pathways may drive inflammatory responses that culminate in preterm labor and fetal inflammation even in the absence of recoverable microorganisms in amniotic fluid cultures.

Longitudinal profiling of inflammatory markers in amniotic fluid (AF), cervico-vaginal fluid, and maternal blood enabled a compartment-specific understanding of host immune response to intrauterine infection. Temporally, AF inflammatory response was the earliest with peak cytokine levels at around 24h post exposure, while the cervico-vaginal inflammatory response had a later onset. Consistent with clinical studies of chorioamnionitis [23], there was minimal inflammation in the maternal blood of NHP even in the face of high AF cytokine levels in the same animal. Longitudinal sampling was also exploited to understand temporal relationship between anakinra administration and prevention of preterm labor. Using mixed effects model with the intervention time as a fixed variable, a clearly separated “flat trajectory” was noted in cervico-vaginal lavage fluid inflammatory markers specifically in the subgroup of anakinra treated animals with no preterm labor (Fig. 4C and Fig. S5B). Interestingly, there was no clear separation of trajectory in any experimental group for inflammatory markers in the AF (Fig. S4B). These divergent responses in the AF vs. cervico-vaginal lavage suggest that inflammation in the cervix may be a key driver of preterm labor in this model. Given that soluble mediators in cervico-vaginal fluid have been proposed as biomarkers [32], our findings extend this concept by demonstrating that cellular infiltration further enhances the utility of cervico-vaginal lavage fluid as a mechanistically informative biomarker of inflammation mediated preterm labor.

*IL6* mRNA expression was consistently elevated in the fetal membranes and cervix of animals that underwent preterm labor, consistent with previous studies identifying IL6 as a robust biomarker of preterm labor [33, 34]. Notably, anakinra reduced *E. coli*–induced *IL6* mRNA expression in both tissues regardless of pregnancy outcome, with the greatest suppression of IL6 observed in animals that did not develop preterm labor. Similarly, cord blood IL6 concentrations were reduced in the no preterm labor subgroup compared with the *E. coli* group. These findings suggest that inhibition of IL1 signaling attenuates local and systemic IL6 responses associated with inflammation-induced preterm birth even when the treatment was begun after the onset of infectious insult. Previous studies from our group demonstrated that infiltrating neutrophils are the primary source of TNF, myeloperoxidase, and PTGS2 during chorioamnionitis [9, 14], whereas IL6 is produced predominantly by amniotic mesenchymal and decidual stromal cells [9, 35]. Indeed, anakinra reduced neutrophilic infiltration both in the fetal membranes and the cervico-vaginal lavage.

In contrast to the cervix and fetal membranes, neutrophil infiltration was not significantly increased in the myometrium following *E. coli* exposure, consistent with our previous observations that myometrial inflammation remains relatively modest during inflammation-induced preterm labor [9]. Correspondingly, anakinra did not significantly reduce *E. coli*–induced increases in uterine cytokine mRNA expression, suggesting that its anti-inflammatory effects are more pronounced in gestational tissues with the highest inflammatory burden during preterm birth.

Inflammatory and endocrine signaling are intimately connected in the pathogenesis of labor [36]. Progesterone, especially via its actions mediated by the PR-B isoform, is essential for maintaining pregnancy, and PR-B signaling exhibits reciprocal crosstalk with inflammatory pathways [24, 25, 37]. We previously reported that intrauterine inflammation, particularly IL1β, upregulates the expression of AKR1C1 in decidua stroma cells, the enzyme that reduces progesterone to its biologically inactive form of 20α-hydroxyprogesterone [16, 38]. Our non-human primate experiments critically extend *in vitro* observations that inhibition of IL1 signaling can partially reverse infection induced activation of AKR1C1 even when anakinra therapy was initiated well after onset of inflammation. Overall, the experiments support the hypothesis that inhibition of progesterone/PR-B signaling in chorio-decidua initiates intrauterine infection induced PTL and inhibition of IL1 signaling partially reverses these adverse effects on withdrawal of progesterone/PR signaling.

Although anakinra exerted anti-inflammatory effects across multiple maternal-fetal compartments when analyzed in the full cohort, these effects were substantially more pronounced in animals that did not develop preterm labor. The basis for this differential response remains unclear. Anakinra concentrations in amniotic fluid and cervico-vaginal fluid did not differ between responders and nonresponders (Fig. S2), arguing against inadequate drug exposure or pharmacokinetic variability as the primary explanation. A possibility is that redundancy in inflammatory pathways may account for IL1 receptor blockade in susceptible animals. Candidate mechanisms include TNF- and IL6–dependent pathways, which have been implicated in the pathogenesis of inflammation-induced preterm labor. Consistent with this possibility, combined inhibition of IL1 and TNF signaling [14, 22, 39, 40], or a broad chemokine suppressing strategy [41, 42], may provide greater therapeutic benefit than targeting IL1 alone. Additional host-specific factors, including genetic or epigenetic variation, may also contribute to differential susceptibility to preterm labor. However, these factors are difficult to experimentally control for in outbred non-human primate studies with limited sample sizes.

Several limitations should be considered. First, although preterm labor incidence (primary outcome), demonstrated a clinically meaningful reduction in effect size with anakinra treatment, this did not reach conventional statistical significance. Second, while anakinra showed substantial efficacy when administered after the onset of inflammation induced by the highly virulent pathogen *E. coli*, the generalizability of these findings to other clinically relevant microorganisms remains unknown. In particular, intra-amniotic infection and inflammation associated with less virulent organisms, including *Ureaplasma* species, are more commonly implicated in spontaneous preterm birth in many human populations [43]. Finally, the study evaluated a relatively short period of infection and treatment (≤7 days), which may not fully recapitulate the prolonged or recurrent inflammatory exposures that may be encountered in clinical settings. Future studies will be required to determine the efficacy of IL1 blockade across diverse microbial etiologies and during more sub-acute forms of intrauterine inflammation.

Anakinra use in pregnancy has been found to be safe [44, 45], and prior studies showed efficacy of IL1 blockade in mouse models of preterm labor [46]. Our study crucially extends these observations to a clinically relevant non-human primate model. The observation that IL1 blockade partially preserves progesterone signaling is significant because the only approved progestin for preterm labor, 17-hydroxyprogesterone caproate, has been withdrawn due to lack of efficacy [47]. These findings support further evaluation of IL1 blockade as an adjunctive therapy for inflammation-associated preterm labor and provide a rationale for combination approaches that simultaneously target inflammatory and progesterone-dependent pathways [48].

## MATHERIALS AND METHODS

### Animals

Normally cycling, adult female Rhesus macaques (*Macaca mulatta*) were timed mated and received an ultrasound guided intra-amniotic (IA) inoculation of either 1 ml LB broth + saline (1:1) solution (Controls n=25) or live uropathogenic *E. coli* at approximately 80-85% gestation (n=21) (Fig. 1A). The *E. coli* was a cystitis clinical isolate derivative (strain UTI89), given at 10^6^ CFU in 1ml [9]. The target for inoculation was gestational day 140 (range 137 – 144 days) where term is 165 days. All the *E. coli*-exposed macaques and five Controls were administered antibiotics 24 hours following the bacterial inoculation. The antibiotic dosing regimen was intramuscular (IM) cefazolin 25 mg/kg twice daily and IM enrofloxacin 5 mg/kg twice daily. Additionally, the macaques received cefazolin 10 mg and enrofloxacin 1 mg once daily via ultrasound guided IA injection. The efficacy of the antibiotic therapy was determined by daily AF and maternal blood cultures from 24 hours following the start of the treatment and in fetal blood cultures at the experimental end point. Of the 21 *E. coli*-exposed animals, ten of the macaques were randomized to receive this antibiotic therapy alone, and eleven were randomized to receive this antibiotic regimen alongside treatment with the IL-1 receptor antagonist (IL-1Ra) Kineret (Anakinra; Sobi) starting 24h till day 5 *after E. coli* inoculation (Fig. 1A). Anakinra was given 100 mg subcutaneous twice daily and 50 mg once daily by intraamniotic route. The length and the echogenicity of the uterine cervix was also assessed daily by ultrasound and was recorded to assess for any progressive changes (Fig. S1).

Body temperatures were monitored by an implanted subcutaneous chip (IPTT-300 implantable programmable temperature transponder, BMDS, Seaford, Delaware, USA). Readings were obtained every 12 hours.

The work reported in this manuscript was conducted during two distinct experimental seasons, separated by 13 months. In the first season 12 macaques (n=6 in each arm – with or without anakinra) received daily treatment until day 4 following *E. coli* inoculation, at which point they underwent surgical delivery and necropsy if preterm labor or preterm birth had not occurred prior. In the second season, 9 macaques (n=4 antibiotics only, and n=5 antibiotics + anakinra) received daily treatment until day 7 following *E. coli* inoculation at which point they underwent surgical delivery and necropsy if preterm labor or preterm birth had not occurred prior. The data from two seasons were combined since the readouts were similar and all preterm labor occurrences were prior to or on d4. Preterm labor status was determined by the veterinary surgeon blinded to the group allocations. Formal evaluation was done at least once daily and more frequently as needed. Although partial changes were seen in some animals, the complete spectrum of preterm labor with assessment of imminent delivery was defined as definite progressive cervical changes, including softening, shortening, and dilatation assessed by ultrasound and verified by manual speculum exam performed by highly trained non-human primate veterinary surgeons blinded to the study group (Fig. S1). Preterm birth was defined as spontaneous vaginal delivery of a live fetus.

The animal clinical characteristics are listed in Tables S1-2.

### *E. coli* culture and cell free DNA *E. coli* detection

*E. coli* colony growth using standard clinical bacteriological methods was reported at different time points in the amniotic fluid, maternal plasma, cord blood plasma, and fetal liver tissue lysate as previously described [9]. Microbial cell free DNA (cfDNA) was determined in the amniotic fluid, maternal plasma, cord blood plasma using next-gen sequencing of cell-free DNA with subsequent analyses using the RUO version of Karius assay to differentially identify metagenomic sequences separately from host DNA [17, 28].

### Biological fluid and tissue collection, enzymatic digestion, and cell count

Maternal plasma, amniotic fluid, and cervico-vaginal fluid were collected daily upon animal sedation [49]. At surgery necropsy, extraplacental membranes chorion-amnion decidua parietalis, myometrium, cervix, umbilical cord blood, fetal liver, and fetal lung were collected within 30 minutes of delivery. For some experiments, fetal membranes, myometrium, cervix, and fetal lung were digested enzymatically, and a cell suspension was created as previously described [14, 40, 49]. Briefly, the tissues were finely minced and enzymatically digested with a cocktail of collagenase A (Roche) and dispase (Life Technologies) for 30 minutes at 37°C followed by DNAsel (Roche) for 30 minutes at 37°C. After filtration, the cells were counted using a trypan blue exclusion test and viability was >90%. Approximately 0.5×10^6^ cells of amniotic fluid were spun and the microscope slides stained with Differential Quick stain Kit. Polymorphonucleated neutrophils counts were performed in a blinded manner.

### Flow cytometry

Monoclonal antibodies (mAbs) used for multiparameter flow cytometry (LSR Fortessa 2, BD Biosciences, San Diego, California, USA) were used to stain cell suspension obtained after enzymatic digestion of fetal membranes, myometrium, cervix, and fetal lung (Table S3) and the gating strategy used to identify the different leukocyte subpopulations was performed as previously described [14]. All antibodies were titrated for optimal detection of positive populations and mean fluorescence intensity. At least 500,000 events were recorded for each sample. Doublets were excluded based on forward scatter properties, and dead cells were excluded using LIVE/DEAD Fixable Aqua Dead Cell Stain (Life Technologies). Unstained and negative biological population were used to determine positive staining for each marker. Following the staining all samples were resuspended in Stabilizing Fixative solution (BD Biosciences) and processed within 30 minutes. Data were analyzed using FlowJo version 9.5.2 software (TreeStar, Ashland, Oregon, USA).

### Quantitative RT-PCR

At delivery, fetal membranes, myometrium, cervix, and fetal lung biopsies were immediately snap-frozen in liquid nitrogen and stored at −80°C. Total RNA was extracted from these biopsies after homogenizing in TRIzol (Invitrogen, Waltham, MA, USA). RNA concentration and quality were measured by NanoDrop spectrophotometer (Thermo Fisher Scientific, Wilmington, Delaware, USA). Reverse transcription of 500 ng RNA and quantitative RT-PCR were performed using qScript One-Step RT-qPCR Kit (Quanta BioSciences, Beverly, MA, USA) following the manufacturer’s instructions and with Rhesus-specific TaqMan gene expression primers (Life Technologies, Table S4). Eukaryotic 18S rRNA (Life Technologies) was endogenous control for normalization of the target RNAs.

### Immunohistochemistry

Formalin fixed, paraffin embedded full thickness fetal membrane parietalis containing amnion, chorion and decidua were sectioned (5 µm mounted on glass slides) subjected to immunostaining as previously described [15, 16, 50]. Sections were first subjected to antigen retrieval at 125°C in using a citric acid-based antigen unmasking solution (Vector Labs cat# H-3300) supplemented with 0.05% Tween-20 at 125°C, and then incubated for 30 minutes at room temperature in blocking solution consisting of 2.5% normal horse serum (Vector Labs cat# S-2000-20) in PBS. Sections were then incubated overnight at 4°C with primary antibody diluted 1:200 in antibody diluent (Cell Signaling Technology cat# 8112L). The primary antibodies were rabbit-anti-PR-B (Cell Signaling Technology cat# #3157) and mouse anti-AKR1C1 (Fitzgerald, cat no. 10R-1776). The following day, sections were washed in tris-buffered (pH 7.4) saline containing 0.05% Tween-20 (TBTS), and then incubated with horseradish peroxidase-conjugated anti-rabbit secondary antibody (Cell Signaling Technology cat# 7074) or horseradish peroxidase-conjugated anti-mouse secondary antibody (Cell Signaling Technology cat# 7076) for 30 minutes at room temperature. After washing in TBST, immunoreactive signal was detected with diaminobenzidine substrate (Cell Signaling Technology cat# 8059) for 3-10 minutes. Sections were stained with hematoxylin, washed in water, dehydrated in increasing concentrations of ethanol and coverslips applied with xylene-based mounting media. Immunoreactive signal was captured by light microscopy.

### Cytokine and chemokine detection

Cytokine/chemokine concentrations in AF, CVL, fetal, and maternal plasma were determined by Luminex detection system employing xMAP magnetic bead technology. Non-human primate specific multiplex kits were used (Millipore, cat# PRCYTOMAG-40K). Values for analytes were extrapolated based on fluorescent intensity curves for standards.

### Anakinra levels in the amniotic fluid and cervico-vaginal lavage fluid

Human IL1ra Luminex detection system (Thermofisher cat# EPX01A-12080-901) was chosen based on a high dynamic range of detection (34.18 pg to 140,000 pg/mL) and higher affinity for human vs. Rhesus macaque IL1ra.

### Statistics

Continuous variables are presented as mean ± SEM. Comparisons among three groups were performed using the Kruskal–Wallis test, followed by Dwass–Steel– Critchlow–Fligner multiple-comparison testing when appropriate. Two-group comparisons were performed using the Wilcoxon signed-rank test or Mann–Whitney U test, as indicated. Categorical variables were compared using Fisher’s exact test or the chi-square test. All tests were two-sided, and P < 0.05 was considered statistically significant. Analyses were performed using SAS version 9.4 (SAS Institute). Because tissue-specific biomarker analyses were mechanistic in nature, P values were not adjusted for multiple comparisons.

Longitudinal cytokine measurements in amniotic fluid and cervico-vaginal lavage fluid were analyzed using linear mixed-effects models with animal-specific random intercepts to account for repeated measures. Fixed effects included treatment group and time. Inflammatory trajectories were modeled as two prespecified epochs: before treatment initiation (0–24 h after *E. coli* inoculation) and after treatment initiation (24 h to delivery). Regression slopes and treatment-by-time interactions were compared across groups.

Given the relatively small sample size, the effect of anakinra on preterm labor incidence was further evaluated using a Bayesian beta-binomial model. Noninformative Beta (1,1) priors were specified for both groups, and posterior distributions were updated using the observed outcomes (*E. coli*, 9/10 animals with preterm labor; *E. coli* + anakinra, 6/11 animals). The posterior distribution of the between-group difference in preterm labor incidence was estimated by Monte Carlo simulation. Results are reported as posterior mean differences, 95% credible intervals, and posterior probabilities of treatment benefit.

## Study approval and ethics statement

All animal procedures were approved by the Institutional Animal Care and Use Committee (IACUC; protocol # 22121) at the University of California Davis and endorsed by the University of California, Los Angeles. The committee that reviewed and approved was the UC Davis IACUC. Care and housing of animals met all IACUC, US Department of Agriculture, and US NIH guidelines for humane macaque husbandry, including the presence of enrichment objects, daily foraging enrichment, and auditory and olfactory access to conspecifics in the same room.

## Acknowledgments

The authors thank Jennifer Kendrick, Sarah Lockwood, Diana Diaz, Paul-Michael Sosa, Naomi Richards, the research personnel at the CNPRC, and Dr. Laura Garzel at University of California Davis, for assistance with the animals and the undergraduate/medical students Anne Vu, McKensie Hammons, Catherine Lirtsman, Alexander Chao, and Rachel Lande for expert technical assistance. We acknowledge the Immune Assessment Core (IAC) for multiplex ELISA at University of California Los Angeles. Fig. 1A was created with BioRender.com (https://BioRender.com).

## Author contributions

P.P., A.H.J, C.A.C., and S.G.K. conceptualized and designed the study. P.P., D.S, M.C., N.P, A.G, A.G., A.DT., J.S., S.B., S.K., P.B. obtained and process animal samples, and perfomed all the experiments;. P.P., L.K, L.A.M, S.S.W., W.J.Z., H.D., S.D., M.S.M, A.H.J, C.A.C, M.R.J, S.M, and S.G.K. participated in analysis and interpretation of data. All authors have reviewed the manuscript and approve the final version.

## References

1. Goldenberg, R.L., J.F. Culhane, J.D. Iams, and R. Romero, Epidemiology and causes of preterm birth. Lancet, 2008. 371: 75–84.

2. DiGiulio, D.B., R. Romero, H.P. Amogan, J.P. Kusanovic, E.M. Bik, F. Gotsch, C.J. Kim, O. Erez, S. Edwin, and D.A. Relman, Microbial prevalence, diversity and abundance in amniotic fluid during preterm labor: a molecular and culture-based investigation. PLoS One, 2008. 3: e3056.

3. Jung, E., R. Romero, L. Yeo, R. Diaz-Primera, J. Marin-Concha, R. Para, A.M. Lopez, P. Pacora, N. Gomez-Lopez, B.H. Yoon, C.J. Kim, S.M. Berry, and C.D. Hsu, The fetal inflammatory response syndrome: the origins of a concept, pathophysiology, diagnosis, and obstetrical implications. Semin Fetal Neonatal Med, 2020. 25: 101146.

4. Cappelletti, M., P. Presicce, and S.G. Kallapur, Immunobiology of Acute Chorioamnionitis. Front Immunol, 2020. 11: 649.

5. Combs, C.A., M. Gravett, T.J. Garite, D.E. Hickok, J. Lapidus, R. Porreco, J. Rael, T. Grove, T.K. Morgan, W. Clewell, H. Miller, D. Luthy, L. Pereira, M. Nageotte, P.A. Robilio, S. Fortunato, H. Simhan, J.K. Baxter, E. Amon, A. Franco, K. Trofatter, K. Heyborne, and N. ProteoGenix/Obstetrix Collaborative Research, Amniotic fluid infection, inflammation, and colonization in preterm labor with intact membranes. Am J Obstet Gynecol, 2014. 210: 125 e121–125 e115.

6. Adams Waldorf, K.M., M.G. Gravett, R.M. McAdams, L.J. Paolella, G.M. Gough, D.J. Carl, A. Bansal, H.D. Liggitt, R.P. Kapur, F.B. Reitz, and C.E. Rubens, Choriodecidual Group B Streptococcal Inoculation Induces Fetal Lung Injury without Intra-Amniotic Infection and Preterm Labor in Macaca nemestrina. PLoS One, 2011. 6: e28972.

7. Romero, R., J. Miranda, T. Chaiworapongsa, S.J. Korzeniewski, P. Chaemsaithong, F. Gotsch, Z. Dong, A.I. Ahmed, B.H. Yoon, S.S. Hassan, C.J. Kim, and L. Yeo, Prevalence and Clinical Significance of Sterile Intra-amniotic Inflammation in Patients with Preterm Labor and Intact Membranes. Am J Reprod Immunol, 2014. 72: 458–474.

8. Lee, A.C., L.C. Mullany, M. Quaiyum, D.K. Mitra, A. Labrique, P. Christian, P. Ahmed, J. Uddin, I. Rafiqullah, S. DasGupta, M. Rahman, E.H. Koumans, S. Ahmed, S.K. Saha, A.H. Baqui, and B. Projahnmo Study Group in, Effect of population-based antenatal screening and treatment of genitourinary tract infections on birth outcomes in Sylhet, Bangladesh (MIST): a cluster-randomised clinical trial. Lancet Glob Health, 2019. 7: e148–e159.

9. Cappelletti, M., P. Presicce, M. Feiyang, P. Senthamaraikannan, L.A. Miller, M. Pellegrini, M.S. Sim, A.H. Jobe, S. Divanovic, S.S. Way, C.A. Chougnet, and S.G. Kallapur, The induction of preterm labor in rhesus macaques is determined by the strength of immune response to intrauterine infection. PLoS Biol, 2021. 19: e3001385.

10. Sadowsky, D.W., K.M. Adams, M.G. Gravett, S.S. Witkin, and M.J. Novy, Preterm labor is induced by intraamniotic infusions of interleukin-1beta and tumor necrosis factor-alpha but not by interleukin-6 or interleukin-8 in a non-human primate model. Am J Obstet Gynecol, 2006. 195: 1578–1589.

11. Romero, R., M. Mazor, and B. Tartakovsky, Systemic administration of interleukin-1 induces preterm parturition in mice. Am J Obstet Gynecol, 1991. 165: 969–971.

12. Hirsch, E., Y. Filipovich, and M. Mahendroo, Signaling via the type I IL-1 and TNF receptors is necessary for bacterially induced preterm labor in a murine model. Am J Obstet Gynecol, 2006. 194: 1334–1340.

13. Dolan, S.M., M.V. Hollegaard, M. Merialdi, A.P. Betran, T. Allen, C. Abelow, J. Nace, B.K. Lin, M.J. Khoury, J.P. Ioannidis, S. Bagade, X. Zheng, R.A. Dubin, L. Bertram, D.R. Velez Edwards, and R. Menon, Synopsis of preterm birth genetic association studies: the preterm birth genetics knowledge base (PTBGene). Public Health Genomics, 2010. 13: 514–523.

14. Presicce, P., C.W. Park, P. Senthamaraikannan, S. Bhattacharyya, C. Jackson, F. Kong, C.M. Rueda, E. DeFranco, L.A. Miller, D.A. Hildeman, N. Salomonis, C.A. Chougnet, A.H. Jobe, and S.G. Kallapur, IL-1 signaling mediates intrauterine inflammation and chorio-decidua neutrophil recruitment and activation. JCI Insight, 2018. 3.

15. Amini, P., D. Michniuk, K. Kuo, L. Yi, Y. Skomorovska-Prokvolit, G.A. Peters, H. Tan, J. Wang, C.J. Malemud, and S. Mesiano, Human Parturition Involves Phosphorylation of Progesterone Receptor-A at Serine-345 in Myometrial Cells. Endocrinology, 2016. 157: 4434–4445.

16. DeTomaso, A., H. Kim, J. Shauh, A. Adulla, S. Zigo, M. Ghoul, P. Presicce, S.G. Kallapur, W. Goodman, T. Tilburgs, S.S. Way, D. Hackney, J. Moore, and S. Mesiano, Progesterone inactivation in decidual stromal cells: A mechanism for inflammation-induced parturition. Proc Natl Acad Sci U S A, 2024. 121: e2400601121.

17. Blauwkamp, T.A., S. Thair, M.J. Rosen, L. Blair, M.S. Lindner, I.D. Vilfan, T. Kawli, F.C. Christians, S. Venkatasubrahmanyam, G.D. Wall, A. Cheung, Z.N. Rogers, G. Meshulam-Simon, L. Huijse, S. Balakrishnan, J.V. Quinn, D. Hollemon, D.K. Hong, M.L. Vaughn, M. Kertesz, S. Bercovici, J.C. Wilber, and S. Yang, Analytical and clinical validation of a microbial cell-free DNA sequencing test for infectious disease. Nat Microbiol, 2019. 4: 663–674.

18. Gomez-Lopez, N., R. Romero, Y. Xu, Y. Leng, V. Garcia-Flores, D. Miller, S.M. Jacques, S.S. Hassan, J. Faro, A. Alsamsam, A. Alhousseini, H. Gomez-Roberts, B. Panaitescu, L. Yeo, and E. Maymon, Are amniotic fluid neutrophils in women with intraamniotic infection and/or inflammation of fetal or maternal origin? Am J Obstet Gynecol, 2017. 217: 693 e691–693 e616.

19. Bukowski, R., Y. Sadovsky, H. Goodarzi, H. Zhang, J.R. Biggio, M. Varner, S. Parry, F. Xiao, S.M. Esplin, W. Andrews, G.R. Saade, J.V. Ilekis, U.M. Reddy, and D.A. Baldwin, Onset of human preterm and term birth is related to unique inflammatory transcriptome profiles at the maternal fetal interface. PeerJ, 2017. 5: e3685.

20. Nadeem, L., R. Balendran, A. Dorogin, S. Mesiano, O. Shynlova, and S.J. Lye, Pro-inflammatory signals induce 20alpha-HSD expression in myometrial cells: A key mechanism for local progesterone withdrawal. J Cell Mol Med, 2021. 25: 6773–6785.

21. Hanley, G.E., S. Munro, D. Greyson, M.M. Gross, V. Hundley, H. Spiby, and P.A. Janssen, Diagnosing onset of labor: a systematic review of definitions in the research literature. BMC Pregnancy Childbirth, 2016. 16: 71.

22. Presicce, P., M. Cappelletti, P. Senthamaraikannan, F. Ma, M. Morselli, C.M. Jackson, S. Mukherjee, L.A. Miller, M. Pellegrini, A.H. Jobe, C.A. Chougnet, and S.G. Kallapur, TNF-Signaling Modulates Neutrophil-Mediated Immunity at the Feto-Maternal Interface During LPS-Induced Intrauterine Inflammation. Front Immunol, 2020. 11: 558.

23. Dulay, A.T., I.A. Buhimschi, G. Zhao, M.O. Bahtiyar, S.F. Thung, M. Cackovic, and C.S. Buhimschi, Compartmentalization of acute phase reactants Interleukin-6, C-Reactive Protein and Procalcitonin as biomarkers of intra-amniotic infection and chorioamnionitis. Cytokine, 2015. 76: 236–243.

24. Zakar, T. and F. Hertelendy, Progesterone withdrawal: key to parturition. Am J Obstet Gynecol, 2007. 196: 289–296.

25. Peters, G.A., L. Yi, Y. Skomorovska-Prokvolit, B. Patel, P. Amini, H. Tan, and S. Mesiano, Inflammatory Stimuli Increase Progesterone Receptor-A Stability and Transrepressive Activity in Myometrial Cells. Endocrinology, 2017. 158: 158–169.

26. Merlino, A., T. Welsh, T. Erdonmez, G. Madsen, T. Zakar, R. Smith, B. Mercer, and S. Mesiano, Nuclear progesterone receptor expression in the human fetal membranes and decidua at term before and after labor. Reprod Sci, 2009. 16: 357–363.

27. Baxter, C., I. Crary, B. Coler, L. Marcell, E.M. Huebner, S. Rutz, and K.M. Adams Waldorf, Addressing a broken drug pipeline for preterm birth: why early preterm birth is an orphan disease. Am J Obstet Gynecol, 2023. 229: 647–655.

28. Presicce, P., D. Beckman, G.B. Diniz, M. Cappelletti, S. Ott, S. Bercovici, S. Kale, P. Babb, J. Mohole, L.S. Richardson, A.K. Kammala, R. Menon, L.A. Miller, E.E. Crouch, A.H. Jobe, S. Divanovic, C.A. Chougnet, S.S. Way, J.H. Morrison, and S.G. Kallapur, Diffuse neuroinflammation and immature neuron loss in fetal Rhesus macaques after short-term intrauterine infection. J Neuroinflammation, 2026. 23: 60.

29. Gomez-Lopez, N., J. Galaz, D. Miller, M. Farias-Jofre, Z. Liu, M. Arenas-Hernandez, V. Garcia-Flores, Z. Shaffer, J.M. Greenberg, K.R. Theis, and R. Romero, The immunobiology of preterm labor and birth: intra-amniotic inflammation or breakdown of maternal-fetal homeostasis. Reproduction, 2022. 164: R11–R45.

30. Couceiro, J., I. Matos, J.J. Mendes, P.V. Baptista, A.R. Fernandes, and A. Quintas, Inflammatory factors, genetic variants, and predisposition for preterm birth. Clin Genet, 2021. 100: 357–367.

31. Dong, M. and K.A. Fitzgerald, DNA-sensing pathways in health, autoinflammatory and autoimmune diseases. Nat Immunol, 2024. 25: 2001–2014.

32. Chan, D., P.R. Bennett, Y.S. Lee, S. Kundu, T.G. Teoh, M. Adan, S. Ahmed, R.G. Brown, A.L. David, H.V. Lewis, B. Gimeno-Molina, J.E. Norman, S.J. Stock, V. Terzidou, P. Kropf, M. Botto, D.A. MacIntyre, and L. Sykes, Microbial-driven preterm labour involves crosstalk between the innate and adaptive immune response. Nat Commun, 2022. 13: 975.

33. Leanos-Miranda, A., A.G. Nolasco-Leanos, R.I. Carrillo-Juarez, C.J. Molina-Perez, I. Isordia-Salas, and K.L. Ramirez-Valenzuela, Interleukin-6 in Amniotic Fluid: A Reliable Marker for Adverse Outcomes in Women in Preterm Labor and Intact Membranes. Fetal Diagn Ther, 2021. 48: 313–320.

34. Norwitz, E.R., J.N. Robinson, and J.R. Challis, The control of labor. N Engl J Med, 1999. 341: 660–666.

35. Presicce, P., C. Roland, P. Senthamaraikannan, M. Cappelletti, M. Hammons, L.A. Miller, A.H. Jobe, C.A. Chougnet, E. DeFranco, and S.G. Kallapur, IL-1 and TNF mediates IL-6 signaling at the maternal-fetal interface during intrauterine inflammation. Front Immunol, 2024. 15: 1416162.

36. Keelan, J.A., Intrauterine inflammatory activation, functional progesterone withdrawal, and the timing of term and preterm birth. J Reprod Immunol, 2018. 125: 89–99.

37. Tan, H., L. Yi, N.S. Rote, W.W. Hurd, and S. Mesiano, Progesterone receptor-A and -B have opposite effects on proinflammatory gene expression in human myometrial cells: implications for progesterone actions in human pregnancy and parturition. J Clin Endocrinol Metab, 2012. 97: E719–730.

38. Nadeem, L., O. Shynlova, E. Matysiak-Zablocki, S. Mesiano, X. Dong, and S. Lye, Molecular evidence of functional progesterone withdrawal in human myometrium. Nat Commun, 2016. 7: 11565.

39. Toth, A., S. Steinmeyer, P. Kannan, J. Gray, C.M. Jackson, S. Mukherjee, M. Demmert, J.R. Sheak, D. Benson, J. Kitzmiller, J.A. Wayman, P. Presicce, C. Cates, R. Rubin, K. Chetal, Y. Du, Y. Miao, M. Gu, M. Guo, V.V. Kalinichenko, S.G. Kallapur, E.R. Miraldi, Y. Xu, D. Swarr, I. Lewkowich, N. Salomonis, L. Miller, J.S. Sucre, J.A. Whitsett, C.A. Chougnet, A.H. Jobe, H. Deshmukh, and W.J. Zacharias, Inflammatory blockade prevents injury to the developing pulmonary gas exchange surface in preterm primates. Sci Transl Med, 2022. 14: eabl8574.

40. Jackson, C.M., M. Demmert, S. Mukherjee, T. Isaacs, R. Thompson, C. Chastain, J. Gray, P. Senthamaraikannan, P. Presicce, K. Chetal, N. Salomonis, L.A. Miller, A.H. Jobe, S.G. Kallapur, W.J. Zacharias, I.P. Lewkowich, H. Deshmukh, and C.A. Chougnet, A potent myeloid response is rapidly activated in the lungs of premature Rhesus macaques exposed to intra-uterine inflammation. Mucosal Immunol, 2022.

41. Ng, P.Y., D.J. Ireland, and J.A. Keelan, Drugs to block cytokine signaling for the prevention and treatment of inflammation-induced preterm birth. Front Immunol, 2015. 6: 166.

42. Shynlova, O., A. Boros-Rausch, T. Farine, K.M. Adams Waldorf, C. Dunk, and S.J. Lye, Decidual Inflammation Drives Chemokine-Mediated Immune Infiltration Contributing to Term Labor. J Immunol, 2021. 207: 2015–2026.

43. Romero, R., P. Pacora, J.P. Kusanovic, E. Jung, B. Panaitescu, E. Maymon, O. Erez, S. Berman, D.R. Bryant, N. Gomez-Lopez, K.R. Theis, G. Bhatti, C.J. Kim, B.H. Yoon, S.S. Hassan, C.D. Hsu, L. Yeo, R. Diaz-Primera, J. Marin-Concha, K. Lannaman, A. Alhousseini, H. Gomez-Roberts, A. Varrey, A. Garcia-Sanchez, and M.T. Gervasi, Clinical chorioamnionitis at term X: microbiology, clinical signs, placental pathology, and neonatal bacteremia - implications for clinical care. J Perinat Med, 2021. 49: 275–298.

44. Brien, M.E., V. Gaudreault, K. Hughes, D.J.L. Hayes, A.E.P. Heazell, and S. Girard, A Systematic Review of the Safety of Blocking the IL-1 System in Human Pregnancy. J Clin Med, 2021. 11.

45. Faure-Bardon, V., D. Beghin, B. Fautrel, M. Latour, C. Vauzelle, C. Reinaud, E. Elefant, B. Coulm, and B. Marin, Exposure during pregnancy to Il-1 targeted therapies and pregnancy, foetal and neonatal outcomes: a study from the French Teratology Information Service. Rheumatology (Oxford), 2025. 64: 6106–6113.

46. Nadeau-Vallee, M., C. Quiniou, J. Palacios, X. Hou, A. Erfani, A. Madaan, M. Sanchez, K. Leimert, A. Boudreault, F. Duhamel, J.C. Rivera, T. Zhu, B. Noueihed, S.A. Robertson, X. Ni, D.M. Olson, W. Lubell, S. Girard, and S. Chemtob, Novel Noncompetitive IL-1 Receptor-Biased Ligand Prevents Infection- and Inflammation-Induced Preterm Birth. J Immunol, 2015. 195: 3402–3415.

47. Aaron, D.G., I.G. Cohen, and E.Y. Adashi, The FDA Struggle to Withdraw Makena: Problems With the Accelerated Approval Process. JAMA, 2022. 328: 2394–2395.

48. Mesiano, S.A., G.A. Peters, P. Amini, R.A. Wilson, G.P. Tochtrop, and F. van Den Akker, Progestin therapy to prevent preterm birth: History and effectiveness of current strategies and development of novel approaches. Placenta, 2019.

49. Short, D., M. Cappelletti, A. Vu, M.R. Johnson, S.G. Kallapur, and P. Presicce, Protocol for isolating cervico-vaginal fluid cells from Macaca mulatta to study immunological and functional changes during pregnancy. STAR Protoc, 2025. 6: 103927.

50. Merlino, A.A., T.N. Welsh, H. Tan, L.J. Yi, V. Cannon, B.M. Mercer, and S. Mesiano, Nuclear progesterone receptors in the human pregnancy myometrium: evidence that parturition involves functional progesterone withdrawal mediated by increased expression of progesterone receptor-A. J Clin Endocrinol Metab, 2007. 92: 1927–1933.

