## Supplementary figures and tables for "IL1 blockade attenuates *E. coli* induced intrauterine inflammation and preterm labor in Rhesus macaques"

**Fig. S1**

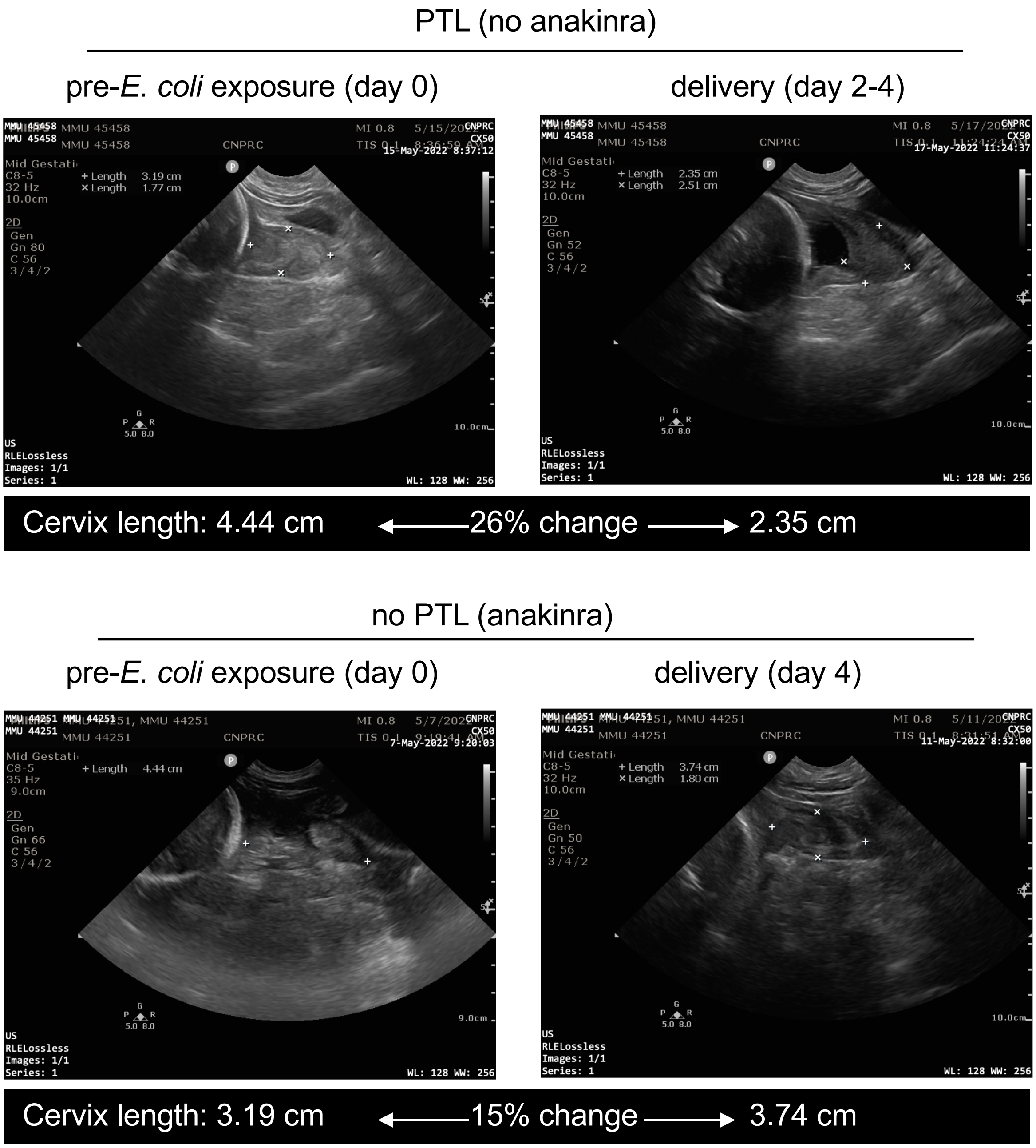


**Fig. S1. Cervical change assessment by ultrasound.** Cervical changes were monitored daily by transabdominal ultrasound to measure cervical length and assess cervical echogenicity, a surrogate marker of inflammation-associated cervical remodeling preceding preterm labor. Definitive assessment of preterm labor were made by manual examination by trained personnel blinded to the study groups. Representative ultrasound images before *E. coli* injection and at delivery showing changes in cervix length in one animal with PTL (top panel) and in one animal treated with anakinra without PTL (bottom). Note the cervix adjacent to fetal cranium is marked by cross marks for length and breadth

**Fig. S2**

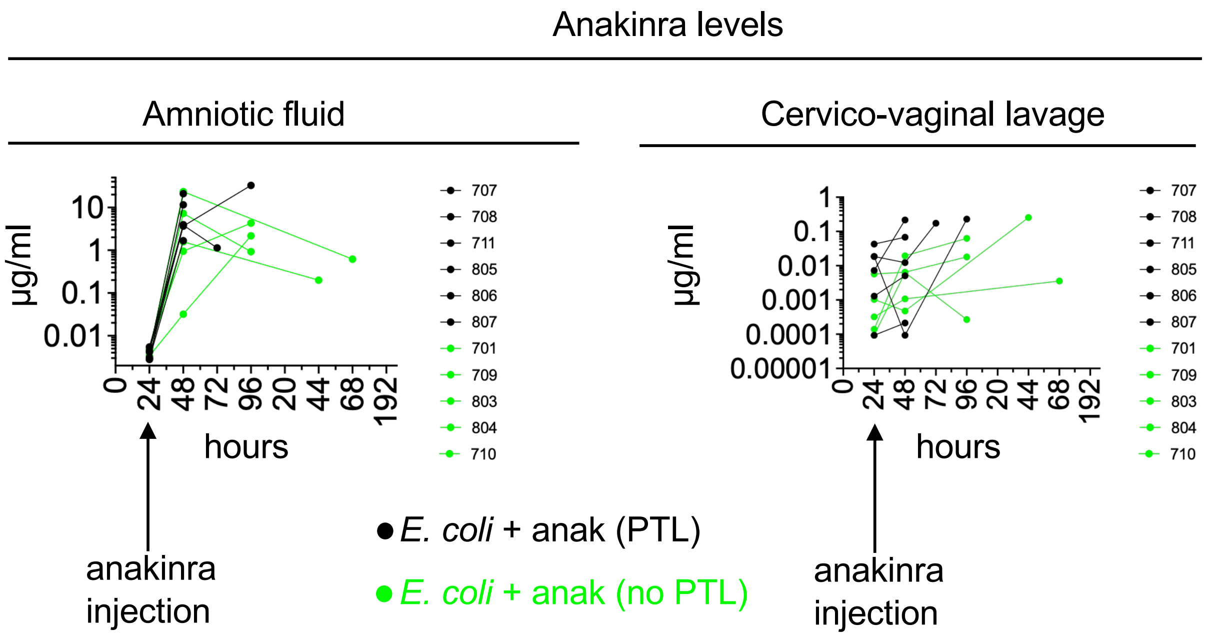


**Fig. S2. Longitudinal levels of recombinant human IL1ra (rhIL1ra) in the amniotic fluid (AF) and cervico-vaginal lavage (CVL).** Recombinant human IL1ra (rh IL1ra) levels were measured were measured since anakinra is a non-glycosated form of rhIL1ra. Luminex bead-based assay was used for longitudinally collected amniotic fluid and CVL samples. Cross-reactivity with rhesus macaque IL1ra was minimized due to higher specificity of the assay for rhIL1ra and much higher drug levels compared to endogenous production allowing use of diluted samples. rhIL1ra level at 24h was prior to the first injection of anakinra in the amniotic fluid. In the amniotic fluid there was a brisk increase in rhIL1ra levels peaking around 48h with a steady decline thereafter. CVL fluid levels of rhIL1ra were much lower compared to the AF and levels increased after drug dosing. In both the AF and CVL, no differences were noted between groups with or without preterm labor.

Fig. S3

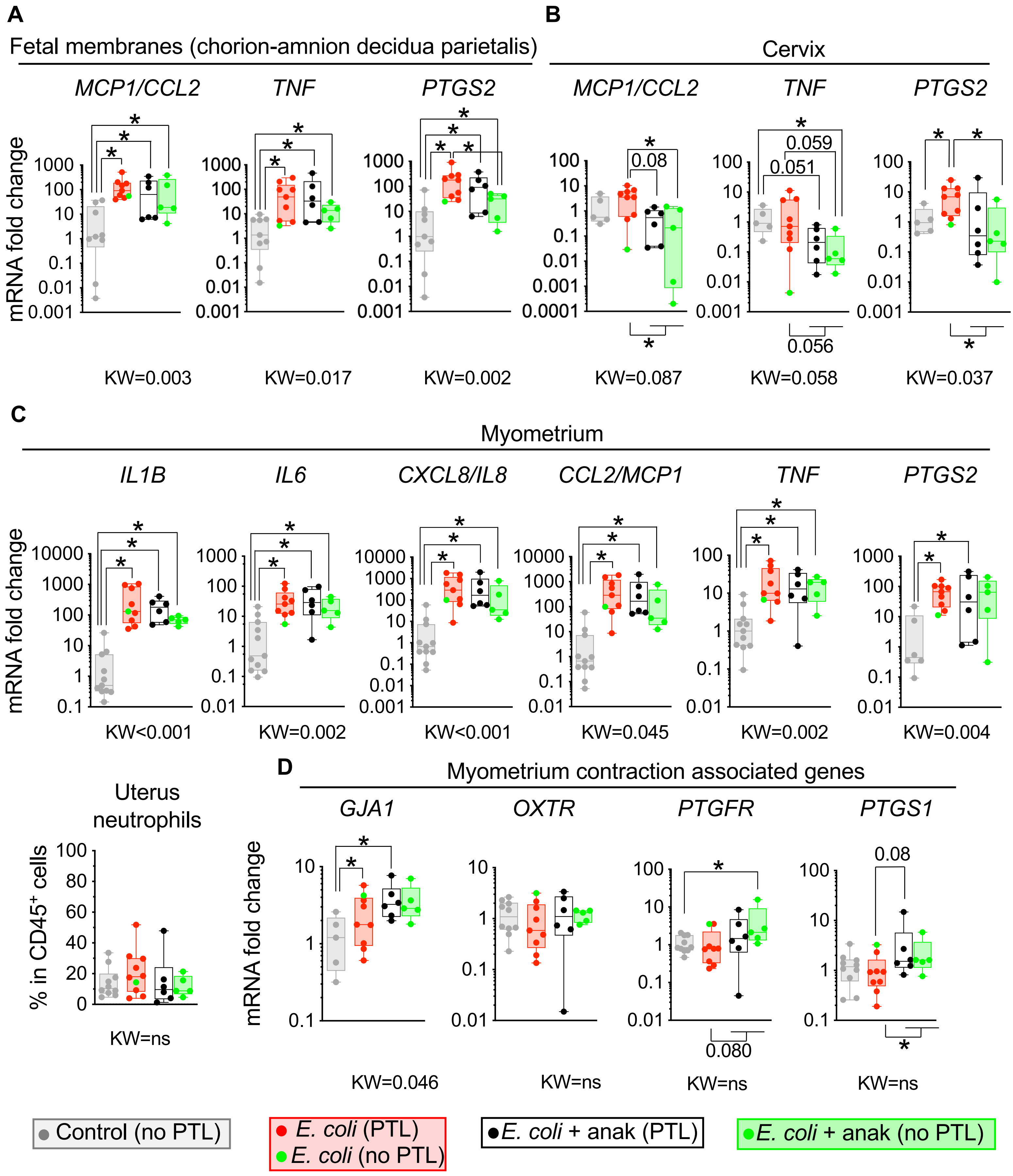

A
Fetal membranes (chorion-amnion decidua parietalis)
MCP1/CCL2
TNF
PTGS2
mRNA fold change
KW=0.003
KW=0.017
KW=0.002
B
Cervix
MCP1/CCL2
TNF
PTGS2
0.08
0.059
0.051
0.056
KW=0.087
KW=0.058
KW=0.037
C
Myometrium
IL1B
IL6
CXCL8/IL8
CCL2/MCP1
TNF
PTGS2
mRNA fold change
KW<0.001
KW=0.002
KW<0.001
KW=0.045
KW=0.002
KW=0.004
Uterus neutrophils
% in CD45+ cells
KW=ns
D
Myometrium contraction associated genes
GJA1
OXTR
PTGFR
PTGS1
mRNA fold change
0.08
0.080
KW=0.046
KW=ns
KW=ns
KW=ns
Control (no PTL)
E. coli (PTL)
E. coli (no PTL)
E. coli + anak (PTL)
E. coli + anak (no PTL)

**Fig. S3. Inflammatory markers in the fetal membranes, cervix, and myometrium at delivery.** Extraplacental chorio-amnion decidua parietalis tissue (fetal membranes), cervix, and myometrium were analyzed for cytokine mRNA expression by quantitative PCR using Taqman probes. Transcript abundance was normalized to 18S RNA and expressed as fold change relative to controls. Neutrophils were quantified by flow cytometry of myometrium cell suspensions. (**A**) *E. coli* exposure significantly increased inflammatory markers in the fetal membranes. Similar results were observed for the anakinra group regardless of PTL. (**B**) In the cervix, *E. coli* exposure increased *PTGS2* mRNA compared to Control. While the decreases in the entire anakinra group were borderline compared to the E.coli group, the anakinra subgroup without PTL had decreased *CCL2/MCP1*, *TNF*, and *PTGS2* expression compared to the *E. coli* group. (**C**) In the myometrium, *E. coli* exposed animals showed a significant increase of *IL1B*, *IL6*, *CXCL8/IL8*, *CCL2/MCP1*, *TNF*, *PTGS2,* mRNAs compared to controls. Similar results were observed for the anakinra group regardless of PTL. There were no differences in neutrophil frequencies between controls and E.coli group with or without anakinra. **(D)** Among the contraction-associated genes, E.coli increased the expression of *GJA1* (connexin-43), but *OXTR, PTGFR,* and *PTGS1* mRNAs did not change. The anakinra subgroup with no PTL had an increased PTGFR mRNA expression compared to the E.coli group. Data are mean ± SEM. Initial all group comparisons were performed using the Kruskal-Wallis (KW) test, followed by Mann–Whitney U-test for two group comparison after stratification of the anakinra group by PTL status. $^*p < 0.05$ between comparators is shown.

**Fig. S4**

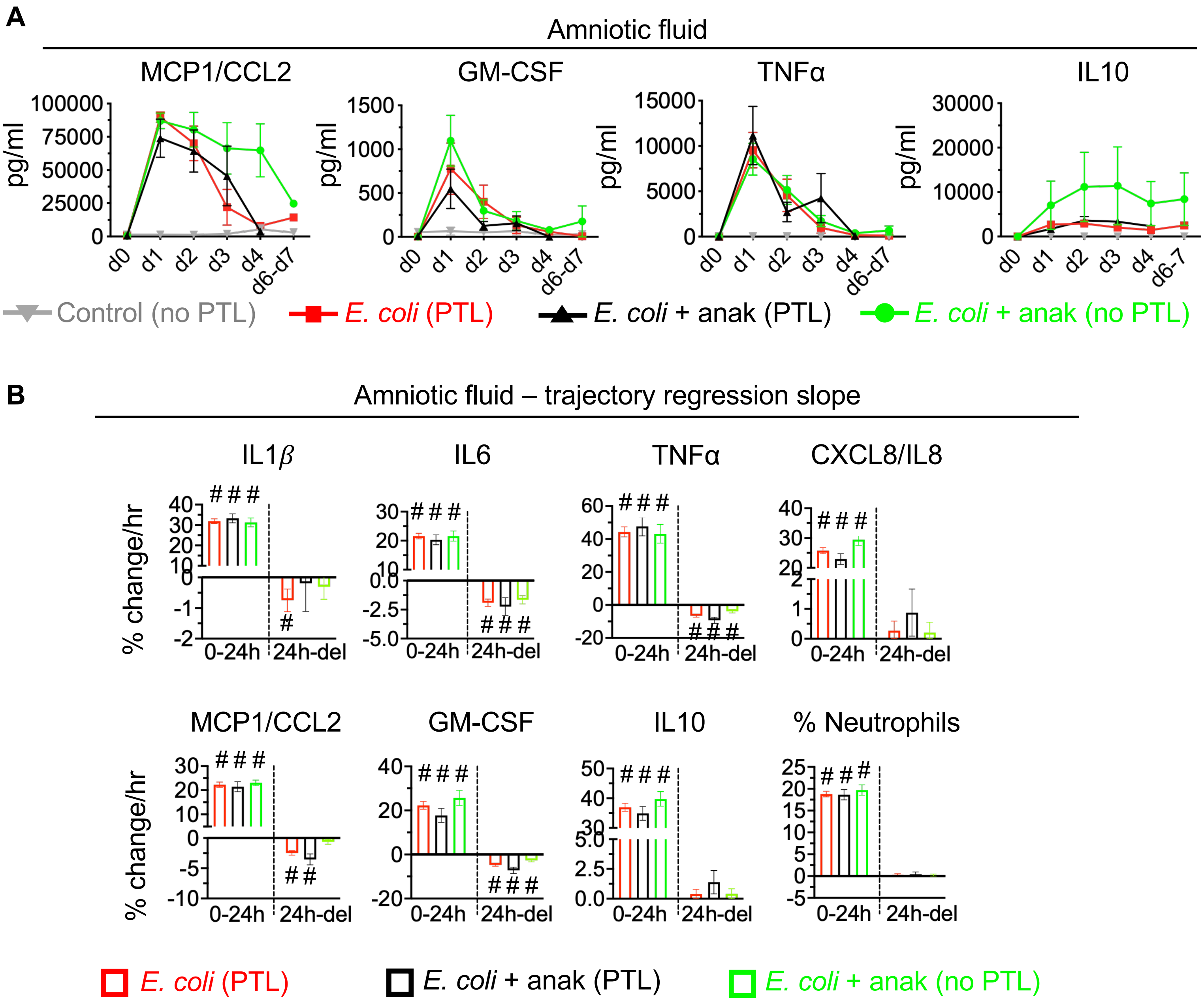


**Fig. S4. Longitudinal profile of amniotic fluid inflammatory markers.** Amniotic fluid was collected longitudinally. **(A)** Cytokine protein concentrations were measured by multiplex assay, and neutrophil frequencies were determined by Diff-Quik staining of cytospin sample. *E. coli*–exposed animals exhibited marked increases in AF pro-inflammatory cytokines with similar temporal kinetics, whereas controls maintained low baseline levels. Temporal kinetics of the anti-inflammatory cytokine IL10 showed sustained increases different from the trajectories of pro-inflammatory cytokines. Neither anakinra treatment nor preterm labor status significantly altered inflammatory marker trajectories. **(B) Regression slope of changes in inflammatory markers in the amniotic fluid.** Mixed-effect model showing regression slopes of cytokine trajectories in the different groups segregated by time periods 0-24h vs 24h to delivery based on treatment intervention at 24h. Note positive slopes in cytokines and neutrophils from 0-24h in all groups denoted by a positive number ($^{\#}p<0.05$ vs a 0 slope). ILB, IL6, MCP1/CCL2, GM-CSF TNF had decreases (negative slopes) from 24h to delivery (del) time period in all groups, while CXCL8/IL8, neutrophils, and IL10 had sustained increases (no change in slope) from the period 24h to delivery.

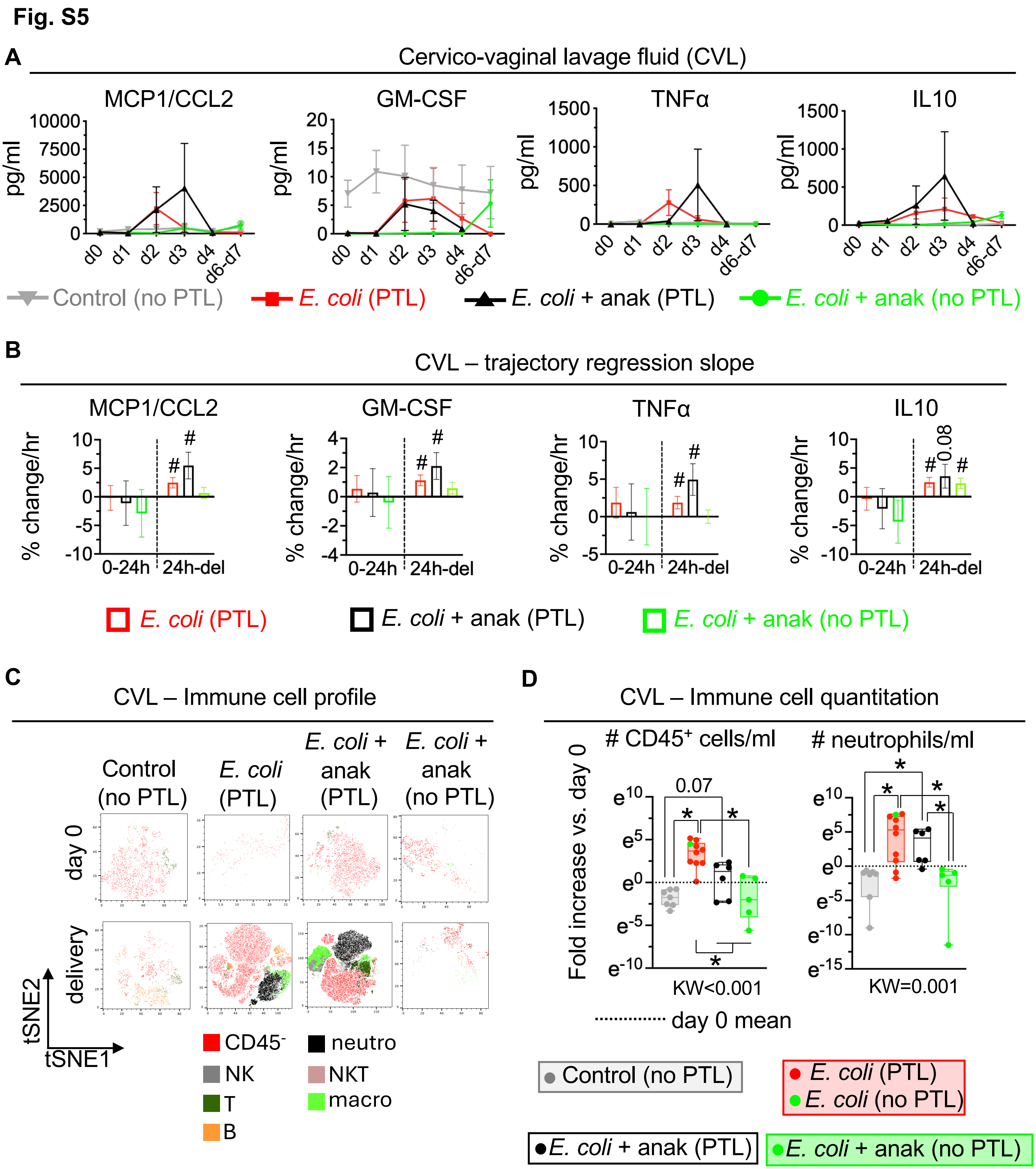

Fig. S5
A
Cervico-vaginal lavage fluid (CVL)
MCP1/CCL2
GM-CSF
TNFα
IL10
pg/ml
d0
d1
d2
d3
d4
d6-d7
Control (no PTL)
E. coli (PTL)
E. coli + anak (PTL)
E. coli + anak (no PTL)
B
CVL – trajectory regression slope
% change/hr
0-24h
24h-del
#
0.08
C
CVL – Immune cell profile
Control (no PTL)
E. coli (PTL)
E. coli + anak (PTL)
E. coli + anak (no PTL)
day 0
delivery
tSNE2
tSNE1
CD45-
neutro
NK
NKT
T
macro
B
D
CVL – Immune cell quantitation
### CD45+ cells/ml
### neutrophils/ml
Fold increase vs. day 0
0.07
*
KW<0.001
KW=0.001
day 0 mean
Control (no PTL)
E. coli (PTL)
E. coli (no PTL)
E. coli + anak (PTL)
E. coli + anak (no PTL)

**Fig. S5. Longitudinal profile of cervico-vaginal lavage (CVL) fluid inflammatory markers.** CVL was collected longitudinally. **(A)** Cytokine protein concentrations were measured by multiplex assay. *E. coli* group and *E.coli* + anakinra subgroup with preterm labor animals increased pro-inflammatory cytokines starting d1 to d2, while the anakinra subgroup with no PTL had a flat trajectory. Controls maintained low baseline levels longitudinally. **(B) Regression slope of changes in inflammatory markers in the CVL.** Mixed-effect model showing regression slopes of cytokine trajectories in the different groups segregated by time periods 0-24h vs 24h to delivery based on treatment intervention at 24h. Note no change in slopes for cytokines for the period 0-24h (slope not different from "0"). For the period 24h to delivery (del), E.coli and E.coli + anakinra subgroup with PTL had increases in MCP1/CCL2, GM-CSF, and TNFα denoted by positive number ($^{\#}p<0.05$ vs a 0 slope), while the anakinra subgroup with no PTL had no increases in these cytokines. For IL10 all groups had a positive slope from the period 24h-delivery ($p=0.08$ for the anakinra subgroup with PTL). **(C)** Representative tSNE plots of flow cytometry data of immune cell populations in the CVL before *E. coli* exposure (d0) and at delivery. Substantial infiltration of different immune cell populations at delivery were noted particularly in the groups experiencing PTL compared to sparse and mostly non-immune cells on day 0. (**D)** $CD45^+$ leukocytes and neutrophil counts at delivery were expressed relative to baseline CVL values in the same animal. *E. coli* markedly increased CD45+ cells and neutrophils at delivery compared to controls. The entire anakinra group had lower CD45+ cells, but only the anakinra sub-group with no PTL had lower neutrophil counts at delivery compared to E.coli. Data are mean ± SEM, Initial all group comparisons were performed using the Kruskal-Wallis (KW) test, followed by Mann–Whitney U-test for two group comparison after stratification of the anakinra group by PTL status. $^{*}p < 0.05$ between comparators is shown.

**Fig. S6**

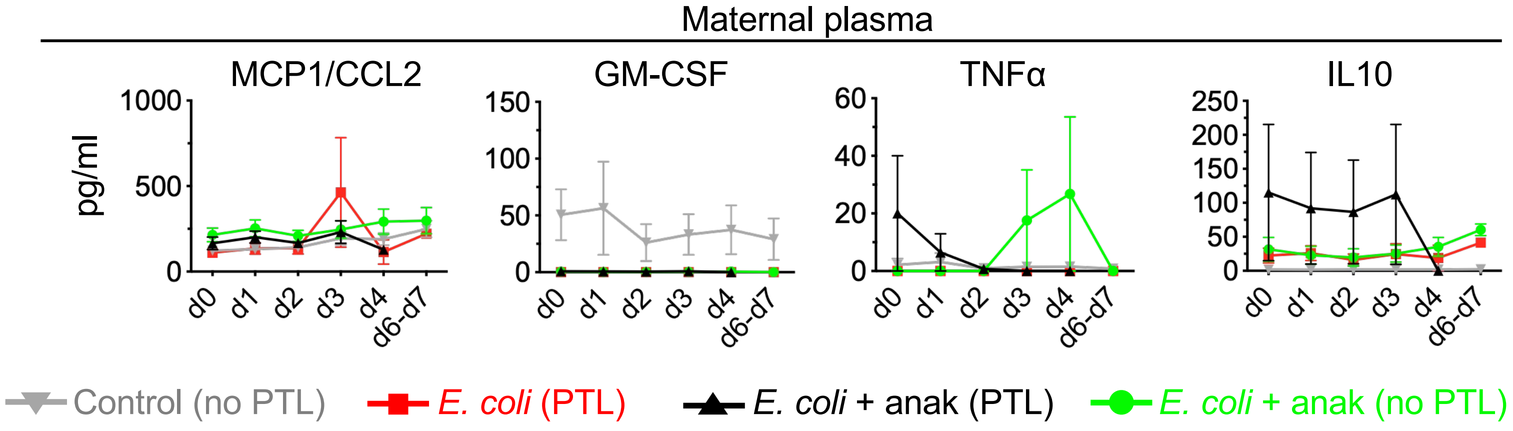


**Fig. S6. Longitudinal profiles of maternal plasma inflammatory markers.** Cytokine protein concentrations were measured by multiplex assay in maternal plasma. All groups including controls had no significant change from baseline values.

**Fig. S7**

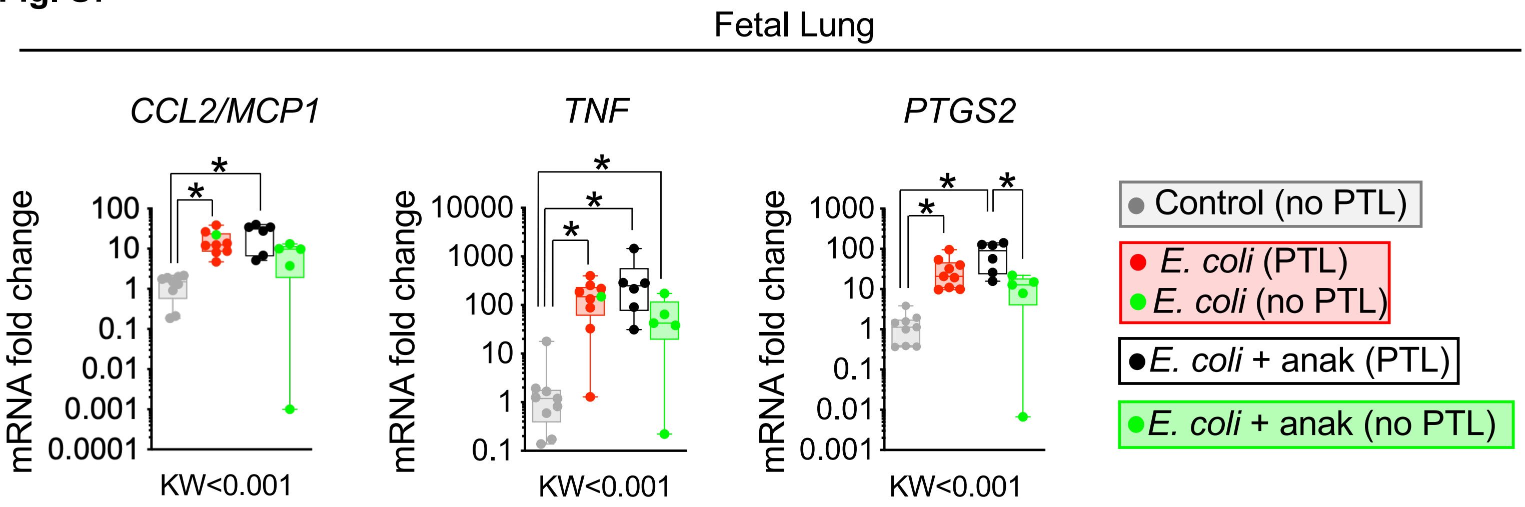


**Fig. S7. Fetal inflammatory markers.** Fetal lung cytokine mRNA expression was evaluated by quantitative PCR using Taqman probes. Transcript abundance was normalized to 18S RNA and expressed as fold change relative to controls. Compared to Control, *E. coli* and anakinra subgroup with PTL increased *CCL2/MCP1, TNF,* and *PTGS2* mRNAs. Compared to Control, anakinra subgroup without PTL increased *TNF* mRNA but not CCL2/MCP1 or PTGS2. Data are mean ± SEM. Initial all group comparisons were performed using the Kruskal-Wallis (KW) test, followed by Mann–Whitney U-test for two group comparison after stratification of the anakinra group by PTL status. *p < 0.05 between comparators is shown.

**Table S1.** Groups of animals and outcome based on preterm labor status

| Rhesus ID | Treatment | Interval between IA *E. coli* injection and delivery (hrs) | PTL/NO PTL | Mode of delivery |
|---|---|---|---|---|
| 700 | *E. coli* + Abx | 91 | PTL | CS |
| 702 | *E. coli* + Abx | 32.5 | PTL | V |
| 703 | *E. coli* + Abx | 58 | PTL | V |
| 706 | *E. coli* + Abx | 78 | PTL | CS |
| 705 | *E. coli* + Abx | 96 | PTL | CS |
| 728 | *E. coli* + Abx | 78 | PTL | CS |
| 800 | *E. coli* + Abx | 70 | PTL | V |
| 801 | *E. coli* + Abx | 16 | PTL | V |
| 802 | *E. coli* + Abx | 72 | PTL | CS |
| 818 | *E. coli* + Abx | 144 | NO PTL | CS |
| 707 | *E. coli* + Abx + anak | 96 | PTL | CS |
| 708 | *E. coli* + Abx + anak | 50 | PTL | CS |
| 711 | *E. coli* + Abx + anak | 52 | PTL | CS |
| 805 | *E. coli* + Abx + anak | 72 | PTL | CS |
| 806 | *E. coli* + Abx + Anak | 50 | PTL | CS |
| 807 | *E. coli* + Abx + Anak | 63.5 | PTL | V |
| 701 | *E. coli* + Abx + Anak | 96 | NO PTL | CS |
| 709 | *E. coli* + Abx + Anak | 96 | NO PTL | CS |
| 710* | *E. coli* + Abx + Anak | 96 | NO PTL | CS |
| 803 | *E. coli* + Abx + Anak | 168 | NO PTL | CS |
| 804 | *E. coli* + Abx + Anak | 144 | NO PTL | CS |
| All Control animals delivered by CS without PTL 96-144 hrs after IA LB broth/saline injection. | | | | |

PTL = preterm labor
CS = C-section
V = Vaginal

**Table S2.** Animal data

| | **Control n=25** | ***E. coli*+ Abx n=10** | ***E. coli* + Abx + Anakinra n=11** |
|---|---|---|---|
| **Maternal age, years (Mean ± SD)** | 11 ± 3 | 8 ± 3 | 10 ± 3 |
| **Maternal body weight in Kg (Mean ± SD)** | 9.6 ± 1.6 | 8.9 ± 2.5 | 10.2 ± 1.7 |
| **Gestational age at delivery, Median [range]** | 135 [129-149] | 145 [140-148] | 145 [140-149] |
| **Neonatal birth weight, gram (Mean ± SD)** | 356.4 ± 52.2 | 382.2 ± 64.3 | 419.1 ± 44.2 |
| **Neonatal gender, Male (M)/Female (F)** | 19M/6F | 7M/3F | 8M/3F |

**Table S3.** List of antibodies used for flow cytometry

| Antibody | Clone | Fluorochrome | Dilution | Company | Catalogue Number |
|---|---|---|---|---|---|
| **Live/Dead** | | Aqua | 1:200 | ThermoFisher | L34957 |
| **CD45** | DO58-1283 | PE-CF594 | 1:10 | BD Biosciences | 562394 |
| **CD3** | SP34-2 | APC- Cy7 | 1:10 | BD Biosciences | 557757 |
| **CD56** | NCAM16.2 | PE-Cy7 | 1:10 | BD Biosciences | 335809 |
| **HLA-DR** | L243 | Brilliant Violet 650 | 1:10 | Biolegend | 307649 |
| **CD88** | P12/1 | Alexa Fluor 647 | 1:10 | Bio Rad | MCA2059 |
| **CD19** | HIB19 | Alexa Fluor 700 | 1:10 | Biolegend | 303325 |
| **CD20** | 2H7 | Alexa Fluor 700 | 1:10 | Biolegend | 302322 |
| **CD123** | 7G3 | FITC | 1:10 | BD Biosciences | 558663 |
| **CD31** | WM59 | PE | 1:10 | Biolegend | 303105 |
| **CD14** | M5E2 | Pacific Blue | 1:10 | Biolegend | 301816 |

**Table S4.** List of probes for qPCR

| Gene target | Vendor | Cat# |
|---|---|---|
| *IL1β* | ThermoFisher | Rh02621711_m1 |
| *IL6* | ThermoFisher | Rh02789322_m1 |
| *CXCL8/IL8* | ThermoFisher | Rh02789781_m1 |
| *MCP1/CCL2* | ThermoFisher | Rh02621753_m1 |
| *TNF* | ThermoFisher | Rh02789783_m1 |
| *PTGS2* | ThermoFisher | Rh02787802_m1 |
| *IL15* | ThermoFisher | Rh02829168_m1 |
| *GJA1* | ThermoFisher | AIY9Y3JRH |
| *OXTR* | ThermoFisher | Rh010141215_m1 |
| *PTGFR* | ThermoFisher | Rh04346483_m1 |
| *PTGS1* | ThermoFisher | Rh00924807_m1 |
